# Functional specialization of the medial temporal lobes in human recognition memory: dissociating effects of hippocampal vs parahippocampal damage

**DOI:** 10.1101/2020.01.25.919423

**Authors:** Georgios P. D. Argyropoulos, Carola Dell’Acqua, Emily Butler, Clare Loane, Adriana Roca-Fernandez, Azhaar Almozel, Nikolas Drummond, Carmen Lage-Martinez, Elisa Cooper, Richard N. Henson, Christopher R. Butler

## Abstract

A central debate in the systems neuroscience of memory concerns whether different medial temporal lobe (MTL) structures support different processes or material-types in recognition memory. We tested a rare patient (Patient MH) with a perirhinal lesion that appeared to spare the hippocampus, using two recognition memory paradigms, each run separately with faces, scenes and words. Replicating reports of a previous case, Patient MH showed impaired familiarity and preserved recollection, relative to controls, with no evidence for any effect of material-type. Moreover, when compared with other amnesic patients, who had hippocampal lesions that appeared to spare the perirhinal cortex, Patient MH showed greater impairment on familiarity and less on recollection, forming a double dissociation. However, when replacing this traditional, binary categorization of patients with a parametric analysis that related memory performance to continuous measures of brain damage across all patients, we found a different pattern: while hippocampal damage predicted recollection, it was parahippocampal instead of perirhinal (or entorhinal) cortex volume that predicted familiarity. Furthermore, there was no evidence that these brain-behavior relationships were moderated by material-type, nor by laterality of damage. Thus, while our data provide the most compelling support yet for dual-process models of recognition memory, in which recollection and familiarity depend on different MTL structures, they suggest that familiarity depends more strongly upon the parahippocampal rather than perirhinal cortex. More generally, our study reinforces the need to go beyond single-case and group studies, and instead examine continuous brain-behavior relationships across larger patient groups.

## Introduction

Ever since the first descriptions of the famous patient HM (Scoville and Milner, 1957), individuals with medial temporal lobe (MTL) damage have been fundamental in delineating the brain regions supporting human memory. Patient studies offer crucial insights into causal brain-behavior relationships, beyond the correlational information afforded by functional imaging in healthy participants. Nevertheless, many questions about the neural basis of human memory remain unresolved (Squire and Wixted, 2011).

In particular, competing accounts have been offered to explain the impact of MTL damage on recognition memory, i.e. the capacity to discriminate whether or not stimuli have been encountered recently. A central question relates to ‘process-specificity’ - whether distinct MTL structures support different processes underlying recognition memory, such as ‘recollection’ (remembering the context in which a stimulus occurred; fundamental for recall) versus ‘familiarity’ (a feeling that a stimulus was encountered, without retrieval of contextual information). According to prominent ‘dual-process’ frameworks (Aggleton and Brown, 1999; Montaldi and Mayes, 2010), recollection relies on the hippocampus (HPC), whereas familiarity relies on regions within the parahippocampal gyrus, principally the perirhinal cortex (PRC). This framework predicts a double dissociation between memory processes, with impaired recollection but not familiarity following selective HPC lesions, and impaired familiarity but not recollection following selective PRC lesions. The main opposing ‘single-process theory’ (Wixted and Squire, 2011a), however, posits that recollection and familiarity reflect subjective expressions of memory traces of varying strength, i.e. familiarity results when weak memory traces fail to re-activate the associated contextual information that characterizes recollection. This theory further proposes that MTL regions function as a single, integrated memory system, such that the degree to which different memory strengths are affected depends on the extent of damage to this MTL system. This theory can explain a single dissociation, whereby lesions can disproportionately affect weak versus strong memories, but not a double dissociation, whereby lesions to different parts of the system have different effects (Montaldi and Mayes, 2010).

However, studies assessing these competing predictions face several challenges: i) standardized neuropsychometric assessment does not suffice to dissociate recollection from familiarity (Argyropoulos and Butler, 2020); ii) there is no universally accepted method of separating recollection and familiarity estimates, so convergent evidence from multiple methods is recommended; iii) selective HPC lesions are rare (Bird and Burgess, 2008), since the conditions associated with these, e.g. ischemia/anoxia, often also cause extra-MTL damage (Huang and Castillo, 2008); iv) selective PRC lesions are even rarer still, with only two cases having been reported: patient IR (Inhoff et al., 2019) and patient NB (Köhler and Martin, 2020). Patient IR had a right PRC lesion and showed perceptual deficits in the absence of memory deficits, though no MTL volumetry was reported, and the memory tasks were not designed to assess familiarity and recollection separately. More importantly, Patient NB did show memory impairments, with impaired familiarity but intact recollection, which is the opposite to the pattern normally reported for HPC lesions, where familiarity is less impaired than recollection. Patient NB’s PRC lesion is therefore vital in providing, in combination with HPC lesions, the double dissociation that favors dual-process theories over single-process theories (Montaldi and Mayes, 2010).

A further challenge for distinguishing dual-versus single-process theories of MTL function concerns v) potential interactions with material-type (Robin et al., 2019). It has been suggested that PRC is important for recognizing objects and faces, while the parahippocampal cortex (PHC), in the posterior parts of parahippocampal gyrus (Pruessner et al., 2002), may be important specifically for scene recognition (Montaldi and Mayes, 2010). Moreover, the entorhinal cortex (ERC) may have specialized routes for object versus scene information, since input from the PRC and the PHC is conveyed into the HPC via different ERC subregions (Maass et al., 2015; Reagh and Yassa, 2014; Van Strien et al., 2009). The HPC has also been claimed to be important for processing scenes (Zeidman et al., 2015), though others propose that its role in memory is independent of material-type (Kim et al., 2015). Material-specific theories provide complementary or even alternative accounts to dual/single-process theories (Lacot et al., 2017), and dissociations previously reported between processes (e.g. recollection vs familiarity) may be specific to certain material-types. Thus, it is important to examine recognition across multiple material-types.

A final challenge for neuropsychological studies concerns vi) the number and definition of patients. Despite the historical influence of single-case studies like HM and NB (Shallice, 2019), testing theories on the basis of single individuals requires consideration of individual differences and measurement noise (Lambon Ralph et al., 2011). Furthermore, defining distinct groups of patients in terms of ‘selective’ lesions to certain brain regions can be misleading (Cipolotti et al., 2006); these groupings are often based on structural brain imaging, from which regions are binarized into “lesioned” or “intact” according to some threshold relative to matched, healthy brains. This categorical approach may miss more subtle, “parametric” relationships between the degree of regional damage and the degree of memory impairment. This latter approach of studying brain-behavior correlations requires larger patient groups, and leverages on individual variability in the distribution of brain damage and memory performance (Argyropoulos et al., 2019).

Here we started by examining an exceptionally rare case of a patient with a lesion in the right PRC (Patient ‘MH’), which appeared to spare the HPC, and who appeared to have memory deficits similar to those reported by Köhler and Martin (2020), i.e, impaired familiarity but intact recollection. In addition to standard neuropsychological tests, we tested him on two paradigms designed to isolate recollection and familiarity in different ways, with the potential to provide convergent evidence. To examine the alternative or moderating factor of material-type, we ran each paradigm with words, unfamiliar faces and unfamiliar scenes. Then, to see how he compared to other patients with MTL damage, we tested another 7 patients on the same paradigms, who, had MRI-confirmed HPC lesions. By comparing these two groups of patients (binarized as either ‘PRC-lesioned/HPC-intact’ or ‘PRC-intact/HPC-lesioned’), we could test for a double dissociation in behavior (as a function of recollection and familiarity, and/or material-type). Going further, we also performed a parametric analysis across all patients between the volumes of the aforementioned regions of interest (ROIs) - i.e. the HPC, PRC, ERC and PHC - and performance on each paradigm, capitalizing on the advantages of this method over categorical, and in particular, single-case, approaches.

## Methods

### Participants

All participants provided written informed consent according to the Declaration of Helsinki. Ethical approval was received from South Central Oxford Research Ethics Committee (REC no: 08/H0606/133).

### Healthy Controls

For the MRI analyses below, patients were compared against a group of 48 healthy controls (CTRs; reported in Argyropoulos et al. (2019); age: median = 64.85; IQR = 15.56 years; sex: 23M:25F). Overall, the patients did not differ from the group of 48 CTRs in terms of M:F ratio (7M:2F; χ^2^=2.71, p = 0.100) or age at research scan (median = 56.93; IQR = 11.78; U = 148, p = 0.142). For the behavioral paradigms, 14 CTRs were recruited through local advertisement (only 6 had available MRI data), 8M:6F, with a mean age at behavioral assessment of 62.11 (SD = 6.20) years, and mean years in education of 13.00 (SD = 1.75). They were all native speakers of English, with no known psychiatric or neurological disorders. Due to scheduling conflicts and technical errors, 12/14 CTR datasets were available for the first paradigm and 9/14 for the second paradigm.

### Patients

All 9 patients (7M:2F; age at behavioral assessment: mean = 60.40; SD = 6.26 years; vs. CTRs: t(21) = 0.66, p = 0.520; education: mean = 12.22, SD = 1.09 years; vs. CTRs: t(21) = 1.19, p = 0.249) were recruited within the context of the Memory and Amnesia Project (https://www.ndcn.ox.ac.uk/research/memory-research-group/projects-1/memory-and-amnesia-project).

#### PRC patient (MH)

This man was 51 years of age at the time of study participation (which did not differ significantly from the mean of CTRs (t(13) = 1.69, p = 0.115) or from the mean of the rest of the patients (t(7) = 1.70, p = 0.132)) and had 12 years of education (which again did not differ significantly from the mean of CTRs (t(13) = 0.55, p = 0.591) or from the mean of the rest of the patients (t(7) = 0.20, p = 0.845)). At the age of 21, while working, he collapsed on the floor, and was hospitalized, where a clinical MRI showed a cerebral abscess in his right PRC, sparing the HPC and other MTL structures. The lesion is illustrated in Figure 1a as a hypointensity in the structural T1-weighted MRI that he underwent at the age of 48 as part of our research study. Volumetric analysis of this MRI (Figure 1b) confirmed that the volume of his right PRC was below a conventional cut-off of Z=−1.67 (i.e. 5^th^ %ile) relative to CTRs (Z=−2.99). This was not true of any of the other MTL ROIs examined, i.e. HPC, ERC, PRC, and PHC. No damage was seen in the left or right amygdala or temporal pole either (all Zs, Z > −1.19). A few years after the incident, he was diagnosed with focal epilepsy, and an EEG disclosed epileptiform activity in the right anterior temporal region. He was treated with the antiepileptic drug carbamazepine and the seizures remitted completely. Neuropsychological assessment (conducted at the age of 48) demonstrated normal levels of intelligence, language, executive function, visuospatial perception, visual and verbal recall, as well as verbal and visual recognition memory (all test scores: Z > −1.67), with the striking exception of recognition memory for faces (Z = −2.33) (Supplementary Table 1).

**Figure 1.**
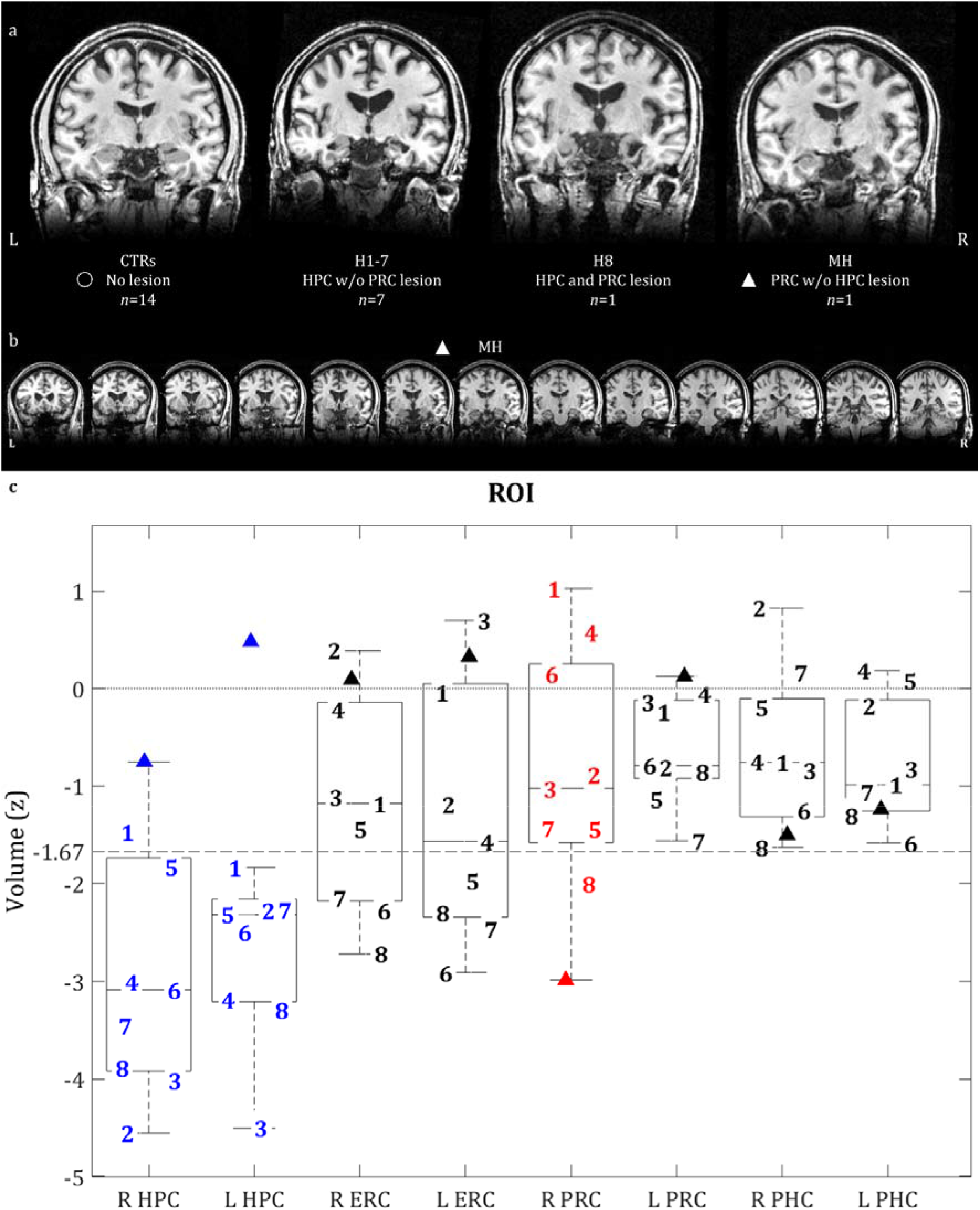
**a:** Coronal slices of structural MR images of CTRs and patients; **b:** A series of coronal slices for MH, highlighting his lesioned right PRC and spared left PRC, along with the rest of his spared MTL structures; **c:** ROI volumes for all 9 patients; boxplots pertain to all 9 patients; line within each boxplot = median value; bottom of box = 25^th^ %ile; top of box = 75^th^ %ile; upper and lower whiskers = scores outside the middle 50; whiskers = 1.5 * interquartile range. **Key: ▲** = MH (patient with right PRC lesion); **1-7** = patients with HPC but no PRC lesion; 8 = patient with both HPC and right PRC lesions; **ERC =** entorhinal cortex; **HPC** = hippocampus; **L, R** = left, right hemisphere; **PHC** = parahippocampal cortex; **PRC** = perirhinal cortex; **ROI** = region of interest (L / R HPC, ERC, PRC, PHC); **Z** = volumes are corrected for TIV and then expressed as Z-scores, based on the mean and standard deviation of the volumes of 48 CTRs whose MTL structures were manually delineated (see Argyropoulos et al., 2019, for details); lesion defined as a Z<-1.67; horizontal lines: Z=0 (dotted line) and Z=−1.67 (dashed line).

#### HPC patients (H1-H8)

We also tested 8 further patients (‘H1-H8’) who all had HPC lesions due to autoimmune limbic encephalitis, which was diagnosed according to consensus criteria (Graus et al., 2016). Volumetric analysis of their T1-weighted MRIs confirmed that all 8 of them had Z-scores below −1.67 relative to CTRs in either left, right, or both HPC (Figure 1b). We call this group the ‘HPC group’. In some of these patients, their ERC volume was also below this cut-off, and for one of the patients (H8), the right PRC was below the cut-off (as well as the left amygdala). We excluded the latter patient for categorical analyses below, but included him in the parametric analysis. None of the remaining MTL ROIs showed volumes below this cut-off.

These patients were representative of the clinical and neuropsychological group-level profile of the autoimmune limbic encephalitis cohort presented in Argyropoulos et al. (2019): i) they were all native speakers of English; ii) they were all recruited after the acute phase of the disease had resolved and were clinically stable (delay from symptom onset range: 1.77-14.92 years); iii) in their acute clinical T2-weighted MRI scans, all 8/8 patients had shown abnormalities in the HPC with respect to volume, T2 signal intensity, and/or diffusion (in one patient, there was also high T2-signal and swelling noted in the right amygdala); in 7/8 patients (H1-H7), these clinically-defined abnormalities did not extend to the parahippocampal gyrus, whereas in the case of one patient (H8), abnormalities had been also noted in both the HPC and the parahippocampal gyrus, and he was the only HPC patient with a lesion in the (right) PRC; iv) no abnormalities were detected beyond the MTL in the research scan that patients underwent acutely or post-acutely (delay from symptom onset range: 1.72-12.93 years); v) in their post-acute neuropsychological assessment (delay from symptom onset range: 1.69-12.93 years), they all showed average to above-average premorbid intelligence [National Adult Reading Test (Nelson and Willison, 1991)], along with vi) preserved (post-morbid) intelligence, semantic memory and language [Wechsler Abbreviated Scales of Intelligence: Vocabulary, Similarities, Matrices (Wechsler, 2011); Graded Naming Test (Mckenna and Warrington, 1980); Camel and Cactus test (Bozeat et al., 2000)]; vii) executive function [Delis-Kaplan Executive Function System - Trails: Number-Letter Switching (Delis et al., 2001)] including working memory [Wechsler Memory Scale: Digit Span forward and backward (Wechsler, 1997a)] (individual impairment on a test was defined as an age-scaled standardized score of ≤-1.67, corresponding to the 5^th^ %ile, in line with standard neuropsychological practice [e.g. (Butler et al., 2014)]), and viii) visuospatial perception [Visual Object and Space Perception battery: cube analysis, dot counting, position discrimination (Warrington and James, 1991)] (all scores above the 5^th^ %ile cut-off point employed in this test); ix) none of the patients had a history of pre-morbid psychiatric or neurological disorder that could have resulted in cognitive impairment and x) none had any contraindication to MRI at the time of entry into the study. Importantly, 7/8 of the HPC patients showed impaired performance in at least one test of anterograde memory [Wechsler Memory Scale (Wechsler, 1997b); Rey-Osterrieth Complex Figure Test (Rey, 1959); the Warrington Recognition Memory Tests for faces and words (Warrington, 1984) and the Warrington Topographical Memory test for scenes (Warrington, 1996); the Doors and People test (Baddeley et al., 1994)], (Supplementary Table 1). Interestingly, only patient H8 (the HPC patient who also had right a PRC lesion) showed impaired face recognition memory (like MH above).

### Scanning procedures

We acquired 3D T1-weighted MRIs for all 9 patients (Siemens 3T Trio system; 32-channel head coil; University of Oxford Centre for Clinical Magnetic Resonance Research) using a Magnetization Prepared Rapid Gradient Echo (MPRAGE) sequence (echo time = 4.7ms; repetition time = 2040ms; flip angle = 8°; field of view = 192mm; voxel size = 1 × 1 × 1mm) for all patients.

### Manual volumetry

Manual segmentation of hippocampal and parahippocampal ROIs (left / right ERC, PRC, and PHC) was conducted in native space (using ITK-SNAP (Yushkevich et al., 2006)) by a trained researcher (ARF) according to segmentation procedures based on published atlases and protocols (Insausti et al., 1998; Pruessner et al., 2002), described in Loane et al. (2019). The volumes of all structures were corrected for (divided by) total intracranial volume (TIV), calculated from the unified segmentation procedure in SPM12.

### Whole-brain Voxel-based Morphometry (modulated grey matter)

In order to ensure that our group of HPC patients (n=8; H1-8) did not present with GM volume reduction beyond the MTL, we conducted a VBM analysis contrasting the wholebrain modulated GM tissue maps (reflecting GM volume) of the HPC patients against those of 67 datasets of CTRs [previously presented in Argyropoulos et al. (2019)]. VBM was conducted using the Statistical Parametric Mapping software (SPM12 version 7219; http://www.fil.ion.ucl.ac.uk/spm/software/spm12) in Matlab R2020a. Images were examined for scanner artefacts and reoriented to have the same point of origin (anterior commissure) and spatial orientation. They were then bias-corrected to remove intensity non-uniformities, and segmented into GM, white matter (WM), and cerebrospinal fluid (CSF) with the unified segmentation procedure. The diffeomorphic anatomical registration through the exponentiated lie algebra (DARTEL) toolbox was applied to participants’ GM, WM, and CSF to refine inter-subject registration, and study-specific GM templates were generated (Ashburner, 2007). After affine registration of the GM DARTEL templates to the tissue probability maps in MNI (Montreal Neurological Institute, Quebec, Canada) space, non-linear warping of GM images was performed to this template in MNI space. Voxel values in the tissue maps were modulated by the Jacobian determinant that was calculated during spatial normalization, with modulated GM images reflecting tissue volume. These images (voxel size: 1 mm^3^ isotropic) were smoothed using a Gaussian filter of 4 mm FWHM. We then compared GM volume between the group of 8 HPC patients and that of 67 CTRs [contrast: ‘CTRs > HPC patients’; second-level between-subject covariates: age, sex, TIV, study (see Argyropoulos et al. (2019) for details)]. We examined peaks surviving whole-brain FWE-correction (p<0.05). Volume reduction was exclusively noted within the MTL (Supplementary Figure 1). Given the effects of registration and smoothing in VBM, we relied on gold-standard manual volumetry to quantify MTL volumes and examine structure-behavior relationships.

### Behavioral Paradigms

Two recognition memory paradigms were employed to dissociate recollection from familiarity, and each paradigm repeated for unfamiliar human faces, unfamiliar natural scenes, and visually presented, high-frequency words.

### ROC

The first paradigm used receiver operating characteristics (ROC), derived from the distribution of confidence responses across studied and unstudied items, for independent estimation of recollection and familiarity [see (Yonelinas, 1994; Yonelinas and Parks, 2007) for methods]; Supplementary Figure 2). This method has been employed in several studies that examine the impact of MTL lesions on recognition memory [e.g. (Aggleton et al., 2005; Bowles et al., 2007; Yonelinas et al., 2002)].

#### Stimulus Materials

##### Faces

160 photos (targets: n=80;foils:n=80) of faces (front view) of unknown Caucasian individuals with a broad age range (18-91 years of age) were taken from the Face Database (Minear and Park, 2004). All photos were taken under natural lighting and had a neutral grey background provided by a portable projection screen. The target and foil faces were matched for age (targets: M = 61.50, IQR = 48.00; foils: M = 58.00, IQR = 45.50) and for M:F ratio (targets: 24:56; foils: 26:54). They were presented in the center of the display.

##### Scenes

160 pictures of unfamiliar natural landscapes (targets: n = 80; foils: n = 80) were chosen from the royalty-free platform Shutterstock (https://www.shutterstock.com), to include no sign of manmade features (buildings, objects), or of people or animals. The scenes used as targets were selected to resemble those used for the foils with respect to their general theme (Autumn: 4:4; Beach: 6:5; Clouds: 4:4; Desert: 7:7; Forest: 5:6; Hills: 7:6; Lake: 5:6; Mountains: 8:7; River: 6:4; Rocks: 10:11; Sea: 4:6; Waterfalls: 4:6; Winter: 10:8). They were presented in the center of the display (17 cm wide, 11 cm tall).

##### Words

160 words (targets: n=80; foils: n = 80) were common, singular nouns. Those used for targets and foils were matched according to i) corpus frequency [SUBTLEXUS word frequency (Brysbaert and New, 2009)] (targets: M = 15.58, IQR = 28.32; foils: M = 15.14; IQR = 15.78 occurrences per million words; targets vs. foils: U = 3039.50, p = 0.585); ii) length (targets: M = 5; IQR = 2; foils: M = 5, IQR = 2; targets vs. foils: U = 3074.50, p = 0.658); iii) mean concreteness ratings (targets: M = 5.81, IQR = 1.54; foils: M = 5.80; IQR = 1.87; targets vs. foils: U = 1684.00, p = 0.631); iv) mean imageability ratings (targets: M = 5.91; IQR = 1.39; foils: M = 5.97, IQR = 1.74; targets vs. foils: U = 1688.50, p = 0.648); v) mean familiarity ratings (targets: M = 5.65, IQR = 0.86; foils: M = 5.71, IQR = 0.77; targets vs. foils: U = 1701.50, p = 0.698); vi) age of acquisition (targets: mean = 3.43; SD = 1.01; foils: mean = 3.34; SD = 0.91; targets vs. foils: t = 0.49, p =0.63); vii) mean ratings of arousal levels (targets: mean = 4.49; SD = 0.94; foils: mean = 4.29; SD = 0.98; targets vs. foils: t = 1.12, p = 0.26); viii) mean valence ratings (targets: M = 5.29; IQR = 0.77; foils: M = 5.20; IQR = 1.06; targets vs. foils: U = 1653.50, p = 0.521) [see (Scott et al., 2018) for details on ratings of concreteness, imageability, familiarity, age of acquisition, arousal, and valence]. They were presented in the center of the display (font size: 28).

##### Procedure

The experiment was written in Matlab, using the Psychophysics Toolbox (v.3) extensions (Brainard, 1997; Kleiner et al., 2007; Pelli, 1997). Each participant was tested in a quiet room. The session lasted approximately 45 minutes. The order of trial blocks is illustrated in Supplementary Figure 2. Each stimulus was first presented to participants in the study phase, before testing recognition memory in the test phase. In both the study and test phases, each trial started with a fixation cross for 0.5 seconds at the center of the display, replaced by a stimulus. In the study phase, participants were asked to judge if each stimulus was “pleasant”, “neutral” or “unpleasant”. Word stimuli were presented for 3 seconds, and face and scene stimuli for 4.5 seconds, irrespective of participants’ response latencies. In the test phase, participants were asked to judge whether the stimulus presented had been previously encountered in the study phase on a 6-point confidence scale (1=definitely new; 2=probably new; 3=maybe new; 4=maybe old; 5=probably old; 6=definitely old), in a self-paced fashion. This phase included all of the stimuli that had been previously presented in the study phase (targets), along with an equal number of novel stimuli (foils). Participants were asked to make full use of the confidence scale. Based upon extensive piloting, we equated levels of difficulty across material-types, using two studies and, correspondingly, two test phases for scenes and faces, but one study and one test phase for words. Moreover, the study phase for words was positioned at the beginning of the session, and the recognition phase for words at the end, similar to other studies [e.g. (Cipolotti et al., 2006)]. The order of blocks and assignment of stimuli to conditions was kept constant across participants, in order to enable the comparison of individual patients with other patients/CTRs. As mentioned above, faces and scenes also remained on the display 1.5 seconds longer than words. Indeed, the parameter estimates (from applying the independent dual-process model to ROCs) for CTRs did not differ across the three material-types for either recollection (one-way repeated-measures ANOVA; independent variable: Material-Type(3); F<0.5, p=0.728; pair-wise t-tests: all ts, |t| ≤ 0.759; all ps, p ≥ 0.464) or familiarity (F<1, p=0.474; pair-wise t-tests: all ts, |t| ≤ 1.23; all ps, p ≥ 0.244), suggesting that the three material-types did not differ with respect to difficulty.

A filler task was also introduced in a series of blocks interspersed within the session, in order to minimize the influence of working memory, as well as to amplify forgetting between study and test phases. In each trial, two numbers were presented side-by-side at the center of the screen. Participants were required to answer a question below those two numbers, asking participants to decide which of the two numbers was higher or lower. Participants selected ‘1’ for the number on the left, ‘2’ for the number on the right, or ‘3’ if the two numbers were equal. Participants were given 3 seconds to respond, before the new trial started.

##### Behavioral Data Analysis

The confidence ratings (ROCs) were analyzed with a dual-process model that assumed recollection and familiarity are independent processes (Yonelinas et al., 1996), using an algorithm available at http://psychology.ucdavis.edu/labs/Yonelinas/DPSDSSE.xls). implemented in Matlab code (http://www.ruhr-unibochum.de/neuropsy/tests/memorysolve.zip), and reported in (Pustina et al., 2012).

### RDP

The second paradigm that was used to provide estimates for recollection and familiarity was based on a response deadline procedure (RDP), which is predicated on the selective reliance of recognition memory on familiarity at short response deadlines, in contrast with long response deadlines (Supplementary Figure 3) (Bowles et al., 2007). The paradigm was administered in two separate sessions, one with a short response deadline (800 ms), and the other with a long response deadline (2,400 ms). The session including the long response deadline was administered first, with a minimum of a 5 days’ delay between the two sessions, so as to prevent interference from the first session in the second session. Patients and CTRs did not differ in the delay between the two sessions (Patients: M = 14; IQR = 227 days; CTRs: M = 14; IQR = 122.50 days; U = 38, p = 0.861). Moreover, we ensured that the first session of the RDP was administered on a different day from the ROC, with a minimum of a 1-day delay across participants. CTRs and patients did not differ on the length of the delay between the ROC and the first RDP session (Patients: M = 30; IQR = 196 days; CTRs: M = 30; IQR = 141.50 days; U = 36, p = 0.712).

#### Stimulus Materials

##### Faces

The 120 faces (n=30 targets and n=30 foils in each deadline condition) were front views of unknown Caucasian people, from the same Face Database as the first paradigm (Minear and Park, 2004), but from different people (see above for more details). The faces used in the short deadline session did not differ from those in the long deadline session in either age (Short Deadline Session: M = 63.00; IQR = 45.75 years of age; Long Deadline Session: M = 61.00; IQR = 47.75 years of age; Short vs. Long Deadline Session: U = 1747.50, p = 0.785) or M:F ratio (Short Deadline Session: 20M:40F; Long Deadline Session: 19M:41F; Short vs. Long Deadline Session: χ^2^ = 0.038, p = 0.845). Targets and foils did not differ with respect to either age or M:F ratio in either the Short (age: targets vs. foils: U = 444.5, p = 0.939; M:F ratio: targets vs. foils: χ^2^ < 0.0005, p > 0.999) or in the Long Deadline Session (age: targets vs. foils: U = 429, p = 0.761; M:F ratio: targets vs. foils: χ^2^ = 0.077, p = 0.781).

##### Scenes

The 120 scenes were pictures of natural landscapes (n = 30 targets and n = 30 foils per deadline condition), taken from the same source as the first paradigm (https://www.shutterstock.com). but different exemplars (see above for details). The Short deadline session did not differ from the Long deadline session with respect to the composition of the different themes across the scenes presented [Short deadline session: autumn(n=3), beach (n=4), clouds (n=3), desert (n=5), forest (n=5), hills (n=5), lakes (n=4), mountains (n=5), rocks (n=7), rivers (n=4), sea (n=5), winter (n=7), waterfalls (n=3); Long deadline session: autumn(n=3), beach (n=5), clouds (n=2), desert (n=5), forest (n=4), hills (n=5), lakes (n=5), mountains (n=4), rocks (n=8), rivers (n=4), sea (n=5), winter (n=7), waterfalls (n=3); Short vs. Long Deadline session: χ^2^ = 0.711, p > 0.999]. Likewise, no such differences were noted between target and foil items in either the Short (χ^2^ = 2.286, p > 0.999) or the Long Deadline session (χ^2^ = 1.810, p > 0.999).

##### Words

The 60 words (n = 30 targets and n = 30 foils) were a subset of those used in the first paradigm (see above for details). Targets and foils did not differ in corpus frequency [SUBTLEXUS word frequency (Brysbaert and New, 2009)] (targets: M = 17.36; IQR = 31.52 occurrences per million words; foils: M = 17.39; IQR= 26.43 occurrences per million words; targets vs. foils: U = 437, p = 0.854); iv) length (targets: M= 5; IQR = 2; foils: M = 5; IQR = 2; targets vs. foils: U = 427, p = 0.741); v) mean concreteness ratings (targets: M = 6.00, IQR = 1.98; foils: M = 5.68; IQR = 1.76; targets vs. foils: U = 252.5, p = 0.995); vi) mean imageability ratings (targets: M = 6.06, IQR = 1.34; foils: M = 6.13, IQR = 1.29; targets vs. foils: U =241, p =0.792); vii) mean familiarity ratings (targets: M = 5.94, IQR = 0.76; foils: M = 5.94, IQR = 0.73; targets vs. foils: U = 239.5, p = 0.766); viii) age of acquisition (targets: M = 2.87; IQR = 1.48; foils: M = 2.60; IQR = 1.43; targets vs. foils: U = 220.5, p = 0.468); ix) mean ratings of arousal levels (targets: mean = 4.54, SD = 1.05; foils: mean = 4.31; SD = 0.94; targets vs. foils: t = 0.77, p = 0.443); x) mean valence ratings (targets: M = 5.47, IQR = 1.04; foils: M = 5.49; IQR = 0.81; targets vs. foils: U = 242.0, p = 0.809).

##### Procedure

Stimulus presentation and data logging were programmed using the Psychophysics Toolbox (v.3) extensions (Brainard, 1997; Kleiner et al., 2007; Pelli, 1997). The session structure is presented in Supplementary Figure 3. The study phase of the paradigm involved 3 blocks (faces, scenes, words) of 30 trials each. Participants were asked to rate each stimulus according to pleasantness (‘Unpleasant’, ‘Neutral’ or ‘Pleasant’). They had 3 seconds to rate pleasantness of words, and 4.5 seconds to rate pleasantness of faces and scenes. In the test phase, participants were required to judge if the item presented on the screen was previously encountered in the study phase (pressing ‘1’ for ‘Old’) or not (pressing ‘9’ for ‘New’). The items were presented over 60 trials, broken down into 6 blocks of 10 trials with breaks after each block.

In each trial, a fixation cross was first presented, followed by the item, which was presented for either 400ms (short response deadline) or 2000 ms (long response deadline). The participant was required to observe the item (face, scene, word) without responding. The item was then bordered in a blue square for 400 ms, during which time the participant was required to provide their response by pressing the ‘OLD’ or the ‘NEW’ button. An error noise was triggered for responses generated before the onset or after the offset of the response window.

For the same reasons as those described for the first paradigm, a series of blocks of filler trials were interspersed within the session, comprising 20 trials each, with a response window of 3 seconds per trial. Participants were presented with two numbers on the screen, and were asked to select which number was the highest or the lowest. They pressed the ‘left’ (arrow to select the number presented on the left side of the display, and the ‘right’ arrow for the number presented on the right side of the display. Participants pressed the ‘down’ arrow to respond that the numbers were equal.

##### Behavioral Data Analysis

Signal detection theory was used to estimate the d’ measure of discriminability for each deadline, which was assumed to reflect pure familiarity in the short deadline, but a combination of recollection and familiarity in the long deadline, such that recollection can be estimated by subtracting the short deadline d’ from the long deadline d’ (Bowles et al., 2007).

### Statistical Analysis

Statistical analysis was performed using R (version 3.5.0) (R Core Team, 2018). The scripts and raw data are available here: https://osf.io/a82ht/

Of the 14 CTRs in total, two had missing values on the first paradigm and five had missing values on the second paradigm. These missing values were imputed using “Multiple Imputation with Chained Equations” (MICE) implemented in the R function “mice”. Five imputations were created, and the results for the analysis below were checked for each imputation (we report the statistics for the first imputation in the main text, but the other four in the Supplementary Tables 2-3).

When comparing patients versus CTRs, we used Analysis of Variance (ANOVA) with within-participant factors Process (Recollection vs. Familiarity), Material-Type (Faces, Scenes, Words), Paradigm (ROC vs. RDP) and a between-participant factor of Group (Patient vs. CTR) using the “aov.res” function in R. Only effects involving the Group factor are of interest and therefore reported in the main Results (for full ANOVA output see Supplementary Tables 2 and 3). In the presence of a significant interaction, lower-order effects are not reported in the main Results, and instead separate, follow-up ANOVAs were performed for each level of one of the interacting factors.

When comparing the two groups of patients against each other (in terms of their Z-scores relative to CTRs, for which imputation was not required), we used a single linear model with the “lmer” function in R, with fixed effects of Group, Paradigm, Process and Material-Type, and a random effect of participant. This model was chosen to make it comparable to the subsequent parametric analysis of brain-behavior relationships across all patients, in which the binary group factor was replaced with continuous measures of Z-scores for the volume of a candidate ROI, plus an additional factor of ROI Laterality. This model was then repeated for the 4 ROIs: HPC, PRC, ERC and PHC. The results of these linear models were reported in terms of Type III Analysis of Variance using Satterthwaite’s method for adjusting degrees of freedom.

## Results

We started with the more traditional categorical analysis of patients, based on a binary classification of whether or not the volume of various MTL structures fell below a threshold (1.67 standard deviations below the mean of the CTR group). More specifically, we examined whether Recollection and/or Familiarity estimates, or the effects of Material-Type, differed i) between the patient with a “selective” PRC lesion (MH) and CTRs, and ii) between a group of patients with a “selective” HPC lesion (n=7, after excluding H8, the HPC case whose right PRC was also below the CTR cut-off) and CTRs. In order to test for a double-dissociation between these two types of patient, we then Z-transformed the patients’ behavioral scores on the basis of the mean and SD of the corresponding CTR data, and examined whether iii) the groups differed in Recollection and/or Familiarity, or the effects of Material-Type. Finally, to examine continuous brain-behavior relationships, we ignored the binary classification into groups, and combined all patients (n=9) for a parametric analysis that related their Z-transformed volumes to their behavioral scores, separately for HPC, PRC, ERC and PHC ROIs. Z-transformed data for patients are shown in Figure 2; raw scores for CTRs and patients are shown in Supplementary Figure 3.

**Figure 2:**
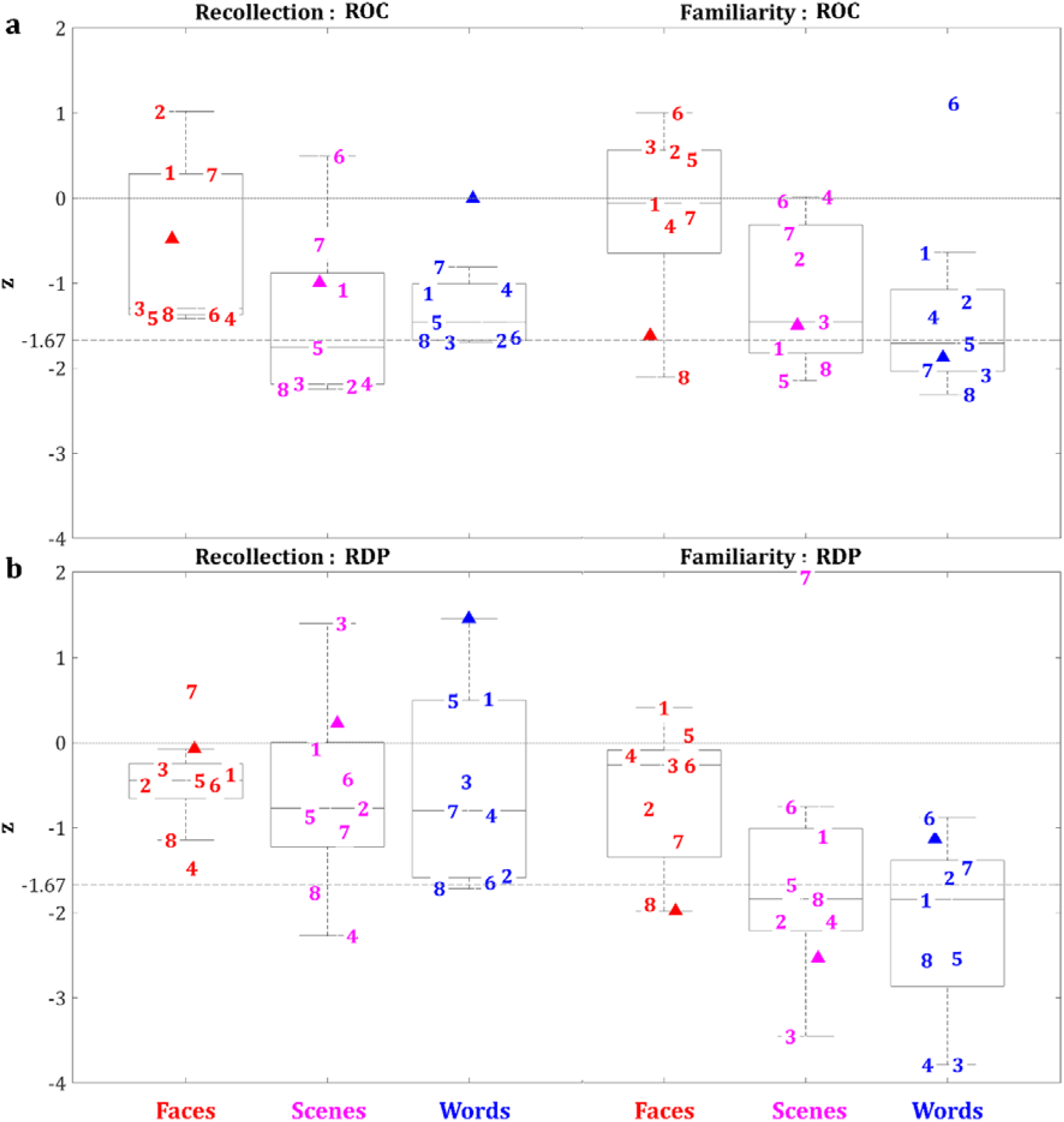
Z-scores for patients’ recollection and familiarity estimates (relative to CTRs) for face, scene, and word recognition memory in the first paradigm (ROC; panel a) and the second paradigm (RDP; panel b). Boxplots pertain to all 9 patients; line in boxplots = median; bottom of box = 25^th^ %ile; top = 75^th^ %ile; whiskers = 1.5 * interquartile range; horizontal lines: Z=0 (dotted line) and Z=-1.67 (dashed line), **key:** □ = MH (patient with right PRC lesion); **1-7** = patients with HPC but no PRC lesion; **8** = patient with both HPC and right PRC lesion.

### MH: single-case study versus CTRs

An ANOVA was conducted on estimates of recollection and familiarity with within-participant factors Process (Recollection vs. Familiarity), Material-Type (Faces, Scenes, Words), Paradigm (ROC vs. RDP) and a between-participant factor of Group (PRC lesion, i.e., MH vs. CTRs). Only effects involving the Group factor are of interest (for full ANOVA output see Supplementary Table 2). The only such effects to reach significance were the main effect of Group (F(1,13) = 7.14, p = 0.019) and the interaction between Group and Process (F(1, 13) = 5.46, p = 0.036). While the latter was only borderline (p = 0.065) for one of the five MICE imputations performed (see Supplementary Table 2), it was consistent for the other four (p < 0.05), so we explored further with follow-up ANOVAs on Recollection and Familiarity separately. These showed no significant effects for Recollection (F < 0.05, p = 0.837), but a significant main effect of Group for Familiarity (F(1,13) =13.74, p = 0.003), with the patient showing lower scores (which was consistent across all imputations). Thus, compared with CTRs, patient MH showed greater impairment for Familiarity than Recollection, consistent with patient NB, with evidence for an impairment on Familiarity but not on Recollection (Bowles et al., 2010, 2007).

### HPC patients: group comparison with CTRs

Unlike the process-specific impairment that we observed above for patient MH, the ANOVA for the HPC group versus CTRs disclosed no such evidence for an interaction between Group and Process (F(1,19) = 1.09, p = 0.310), despite a clear main effect of Group (F(1,19) = 31.42, p < 0.001), where the HPC group performed worse overall (as expected). Follow-up ANOVAs for Recollection and Familiarity both showed a main effect of Group (Recollection: F(1,19) = 7.89, p = 0.011; Familiarity: F(1,19) = 20.14, p < 0.001). Further analyses are reported in Supplementary Table 3.

### Categorical analysis: comparing the two patient groups

To test whether there was a double dissociation between the two categories of patient (PRC vs HPC lesion), we added both MH and the 7 HPC cases (H1-7) to a single linear model, which was fit to their Z-scores relative to CTRs. This model included fixed effects of patient Group (PRC lesion vs. HPC lesion), Paradigm (ROC vs. RDP), Process (Recollection vs. Familiarity) and Material-Type (Faces, Scenes, Words). The full results are shown in Supplementary Table 4, but here we only report significant effects that involve the Group factor, since these reflect dissociations between the behavioral consequences of the two lesion locations. There were two such effects: the two-way interaction between Group and Process (F(1,66) = 8.52, p = 0.005), and the two-way interaction between Group and Material-Type (F(2,66) = 3.20, p = 0.047).

To explore the interaction between Group and Process, we averaged across Material-Type and Paradigm: the mean Z-score for Recollection was −0.81 for the HPC group, but 0.02 for MH, whereas the mean Z-score for Familiarity was −0.94 for the HPC group, but − 1.04 for MH. Although the simple effect of Group did not reach significance for either Recollection (F(1,6)= 3.62, p = 0.106) or Familiarity (F(1,6) = 1.84, p = 0.224) alone, the pattern of means is consistent with a cross-over interaction in the degree of behavioral impairment, with MH relatively more impaired in Familiarity, and the HPC group relatively more impaired in Recollection (consistent with Bowles et al., 2010).

To explore the interaction between Group and Material-Type, we averaged across Process and Paradigm. The mean Z-score for Faces was −0.25 for the HPC group, but − 1.04 for MH, a difference that approached significance (F(1,6) = 5.63, p = 0.055). For Scenes, the mean Z-score was −1.05 for the HPC group and −1.20 for MH, which did not approach significance (F(1,6) = 0.05, p= 0.838). For Words, the mean Z-score was −1.34 for the HPC group and −0.39 for MH, which approached significance (F(1,6)=3.60, p=0.107). Thus, there was some suggestion that MH showed relatively more impairment of overall recognition memory for Faces than the HPC group, and possibly relatively less impairment for Words, but no evidence of a difference for Scenes. Any such effects of Material-Type were explored further in the parametric analysis.

### Parametric Analysis of brain-behavior relationships across all patients

Rather than binarizing patients according to whether specific brain regions are “lesioned” or “intact”, a potentially more powerful way to test models of functional specialization within the MTL is to correlate memory scores with continuous measures of the volume of ROIs (normalized by CTRs), across all patients. For this analysis, we combined MH with all 8 “HPC” patients (H1-H8; including now H8, the patient with right PRC lesion). We fit a single linear model for each of four ROIs: HPC, PRC, ERC and PHC. In each model, the factors of Process, Material-Type and Paradigm were identical to the categorical model above, but the binary Group factor was replaced with the continuous Z-scored volumes of the ROI being tested. We also added a factor for Laterality, in order to capture potential lateralization of function across left and right ROIs. The full results are shown in Supplementary Table 5, but as for the categorical analysis above, here we only report significant effects that involve the ROI volume factor, since these reflect brain-behavior relationships.

For HPC, the only significant effect of interest was the interaction between Volume and Process (F(1,160.33)=17.93, p < 0.001). As shown in Figure 3, when averaging over Material-Type, Paradigm and Laterality, there was a significant positive relationship between HPC volume and Recollection (Figure 3a), but no such relationship for Familiarity (Figure 3b).

**Figure 3:**
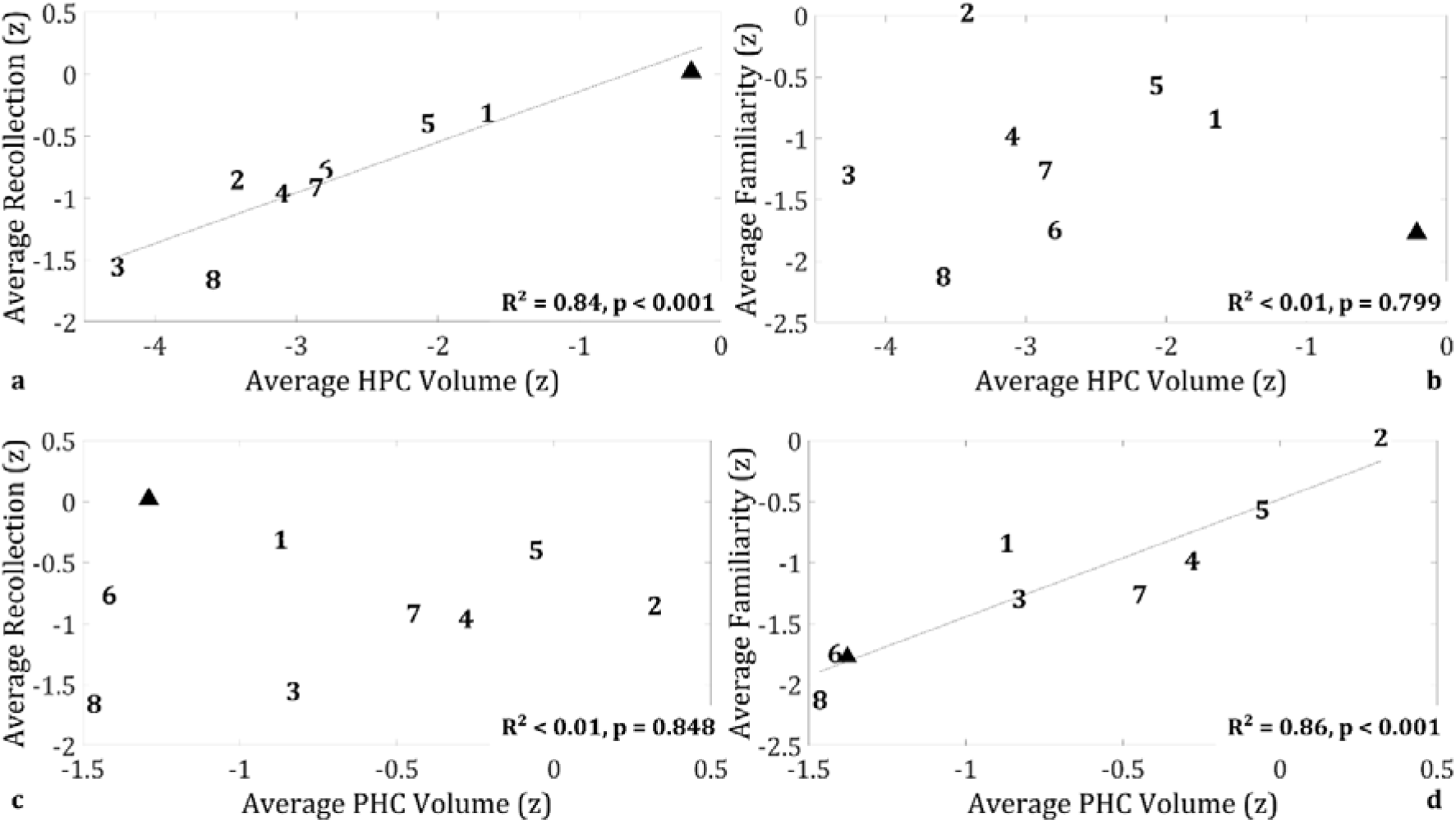
Double dissociation in brain-behavior relationships between HPC volume – Recollection and PHC volume – Familiarity across patients; **a, b:** average HPC volume correlated with Recollection, but not Familiarity; **c, d:** average PHC volume correlated with Familiarity, but not Recollection; **key: HPC:** hippocampus; **PHC:** parahippocampal cortex; **Z:** volumes are expressed as Z-scores, based on the mean and standard deviation of the volumes of the 48 CTRs whose MTL structures were manually delineated (see Argyropoulos et al. (2019) for details); Familiarity and Recollection estimates are expressed as Z-scores, based on the mean and standard deviation of the CTRs that completed the two tasks.

For PRC, no effects involving Volume reached significance, with the interaction with Process having an F-value below 0.5 (see Supplementary Table 5; Supplementary Figure 5). This is a different conclusion from the categorical analysis reported above, and suggests that it is not PRC per se that is important for explaining differences in memory Process or Material-Type.

Likewise, for ERC, there were no significant effects involving Volume (see Supplementary Figure 5; Supplementary Table 5).

However, for PHC, there was a significant interaction between Volume and Process (F(1,159.08)=16.435, p < 0.001). When averaging over Paradigm, Material-Type, and Laterality, there was a significant positive relationship between Volume and Familiarity (Figure 3d), but no such relationship for Recollection (Figure 3c). In other words, the PHC showed the opposite brain-behavior relationship compared with HPC, constituting another double dissociation.

Indeed, if we included both HPC and PHC in a single model predicting Recollection, there was a significant effect of HPC (F(1,14)=71.15, p < 0.001), but not of PHC, nor interaction between HPC and PHC (Fs < 2.30, ps > 0.152). Likewise, if we included both ROIs in a single model predicting Familiarity, there was a significant effect of PHC (F(1,14) = 32.03, p < 0.001) but not of HPC, nor interaction between HPC and PHC (Fs < 1.34, ps > 0.267). These analyses show that HPC makes a unique contribution to Recollection and likewise that PHC makes a unique contribution to Familiarity.

## Discussion

In this study, we examined the relationship between MTL damage and impairment in recognition memory for three different types of memoranda (unknown faces, topographical scenes, and high-frequency words) within two distinct paradigms designed to separate recollection and familiarity processes. Across a cohort of 9 patients with MTL damage, we observed that familiarity impairment was selectively associated with PHC volume reduction, whereas recollection impairment was selectively associated with HPC damage. Our findings provide compelling support for: i) dual-process models of recognition memory, whereby recollection and familiarity depend on the integrity of distinct MTL structures; ii) the PHC as the necessary region for familiarity; iii) the need for future studies to move beyond the case-study approach. We examine each of these three aspects in this order below.

### Support for dual-process models

There is an ongoing debate whether different structures within the MTL support distinct processes underlying recognition memory, especially whether the HPC selectively supports recollection and whether the PRC and PHC selectively support familiarity [e.g. (Aggleton and Brown, 1999; Montaldi and Mayes, 2011); but see (Squire et al., 2004; Wixted and Squire, 2011b)]. Whereas task-based fMRI [e.g. (Kafkas et al., 2016; Staresina et al., 2013a, 2012)] and intracranial EEG studies [e.g. (Staresina et al., 2013b)] have provided some evidence for such dissociations, patient studies are required to establish causal brain-behavior relationships. Nevertheless, patient studies are limited by the scarcity of patients with ‘focal’ PRC lesions. To our knowledge, only one such study has provided evidence for a double dissociation between recollection and familiarity (Bowles et al., 2010). This study reported on patient NB, who had a lesion in the left PRC that extended to the amygdala, ERC and anterolateral temporal cortices, and who presented with a selective familiarity impairment (Bowles et al., 2007). Importantly, she showed the opposite pattern to an older patient who had undergone left amygdalo-hippocampectomy and who presented with a selective recollection impairment (demonstrated with a ‘Remember-Know’ paradigm), despite their comparable overall recognition memory performance (Bowles et al., 2010).

Our PRC patient, MH, has a much more focal lesion in his right PRC than patient NB. Consistent with NB’s performance (Köhler and Martin, 2020), our patient showed impaired familiarity and preserved recollection relative to CTRs. Moreover, when compared directly with a larger group of amnesic patients, who have HPC lesions appearing to spare the PRC, MH showed greater impairment on familiarity and less on recollection, reinforcing a double dissociation. These results are consistent with ‘dual-process’ frameworks [e.g. (Montaldi and Mayes, 2010)]

However, in apparent conflict with dual-process theories and previous reports of impaired recollection but intact familiarity following HPC lesions [e.g. (Aggleton et al., 2005; Vann et al., 2009)], our HPC group did show impaired familiarity relative to CTRs. Prima facie, this finding might be considered to support single-process theories [e.g. (Wixted and Squire, 2011a)], aligning with other reported cases in which HPC lesions are accompanied by impairment of both recollection and familiarity [e.g. (Kirwan et al., 2010; Song et al., 2011; Wais et al., 2006)]. Nonetheless, the latter interpretation was questioned by our subsequent analyses: Moving beyond the dichotomization of structures into ‘lesioned’ and ‘preserved’, we examined whether the variability in familiarity and recollection strength was a function of the volume of different MTL structures across patients. Here, there was compelling evidence for a selective role of HPC in recollection, since HPC volume correlated with recollection but not with familiarity. The impairments in familiarity in our “HPC group” therefore most likely owe to other, subthreshold damage to regions outside the HPC, most likely other regions in parahippocampal cortex (see below). Indeed, the correlation between HPC volume and recollection was mirrored by a correlation between PHC volume and familiarity (but not of PHC volume with recollection). We think this parametric, double-dissociation provides the most compelling evidence to date in favor of ‘dual-process’ theories, particularly those that map these processes to distinct regions with the MTL (e.g. (Montaldi and Mayes, 2010)).

### Possible effects of material-type

Some of our findings may be seen as supporting the view that different parahippocampal structures support familiarity for different material-types (Kafkas et al., 2016; Köhler and Martin, 2020). In particular, both MH and H8 (who had a lesion in the right PRC: volume: Z < −1.67) showed impaired face recognition memory in neuropsychological assessment, along with impaired face familiarity relative to the rest of the patients. At group level, patients with HPC lesions (H1-7) showed only marginal impairment in recollection or familiarity for faces, in contrast with marked impairment for scenes and words (Supplementary Table 3). This pattern dovetails with evidence from case studies of patients with HPC damage [e.g. (Cipolotti et al., 2006)], meta-analytical findings (Bird and Burgess, 2008) on HPC patients’ performance in neuropsychological tests of recognition memory for faces, and with our previous findings on a larger cohort of HPC patients, who showed group-level sparing of face recognition (Argyropoulos et al., 2019). It also supports the idea that the PRC is engaged in processing faces (see (Robin et al., 2019) for discussion). Nevertheless, MH’s familiarity impairment was not selective for faces, and he did not show process-independent but material-specific impairment relative to CTRs. Likewise, our parametric analyses did not detect any evidence for a privileged role of the PHC (e.g. (Buffalo et al., 2006; Litman et al., 2009)) or the HPC (e.g. (Mullally et al., 2014)) in processing spatial information (here, in the form of unknown topographical scenes). Thus, unlike our clear evidence for dissociations between MTL regions as a function of memory process, our results do not fully disentangle potential further dissociations as a function of material-type. Future research with larger patient cohorts (and hence greater power for detecting parametric relationships) may be needed to investigate interactions between memory processes, material-types and MTL regions.

### PHC and familiarity

Within the context of neuropsychological dual-process frameworks, the PRC has been the portion of the parahippocampal gyrus that has been consistently associated with familiarity processes (e.g. (Brown and Aggleton, 2001; Köhler and Martin, 2020)). Less attention has been given to the PHC. In some models, the PHC is considered to support ‘context familiarity’ (e.g. (Montaldi and Mayes, 2010), whereas in others, the PHC is assigned a role in processing context information, including its recollection [e.g. the ‘Binding in Context’ model and subsequent developments (Diana et al., 2007; Ranganath, 2010)]. Our findings here are difficult to reconcile with these models. However, since these are primarily based on task-based fMRI studies of healthy adults, it is possible that PHC contributes to both recollection and familiarity in the healthy brain, but is only *indispensable* for familiarity. In other words, the role of these regions in the healthy brain (disclosed by fMRI) may differ from their role in a damaged brain, owing for example to disruption of connectivity between brain regions (Henson et al., 2016).

### Beyond the case-study approach

Another important methodological message from our findings is the benefit of moving beyond the case-study approach, which requires the dichotomization of structures into ‘lesioned’ (e.g. volumes below some threshold) and ‘preserved’. By testing a larger group, we capitalized on the variability in the integrity of the different structures to examine continuous brain-behavior relationships, allowing for individual differences and obviating the need for arbitrary thresholds to define ‘lesioned vs. non-lesioned’ (Lambon Ralph et al., 2011). The danger of the dichotomous approach to characterizing MTL lesions is illustrated by our examination of the HPC group, who showed evidence of familiarity impairments even though there was no evidence that HPC volume correlated with familiarity. This highlights the possibility that their HPC lesion caused their recollection impairment, but that sub-threshold damage to other MTL regions (e.g. PHC) caused their familiarity impairment. Group-based, parametric analyses like those performed here might help resolve debates in the domain of MTL amnesia that may have arisen largely from the focus on single-case studies [e.g. (Aggleton and Brown, 1999; Montaldi and Mayes, 2011); but see (Squire et al., 2004; Wixted and Squire, 2011b)].

### Limitations and future directions

There are certainly limitations to our study. Firstly, in both paradigms, the order of presentation of the stimulus blocks (faces, scenes, words) was kept constant across participants. This enabled us to compare directly the performance of MH with that of CTRs and HPC patients (since a single-case like MH can typically only attempt one order of tasks, at least when it is difficult to repeat those tasks on the same person; another advantage of group studies is that task order is more easily counter-balanced). Moreover, due to technical errors in the design of the second paradigm, the word stimuli in the second session (long response deadline) were the same as those used in the first session (short response deadline), and were a subset of those used in the first paradigm. Another issue that our study did not address is the possibility that the extent to which scenes are processed by the HPC, PHC, or PRC is a function of stimulus size. It has been argued that relatively small stimuli may be treated like objects, thus maximizing the involvement of the PRC, whereas processing larger background stimuli require an intact PHC (Cassaday and Rawlins, 1997; Montaldi and Mayes, 2010). We thus cannot exclude the possibility that MH’s recognition memory for scenes was affected by the relatively small size of our scene stimuli. These limitations are another reason why the present study cannot offer definitive evidence on the role of different material-types.

Moreover, the nature of our cohort means that we had little power to detect differential effects of bilateral vs. unilateral MTL damage on recognition memory. This is primarily because all 8 of our HPC patients showed largely bilateral HPC damage (L and R HPC: z < −1.4), which is characteristic of the etiology (Argyropoulos et al., 2019). Indeed, bilateral HPC damage may be necessary to see recollection deficits [see (Spiers et al., 2001) for discussion]. Interestingly however, while Patient MH’s PRC lesion was clearly unilateral (R PRC: z = −2.99; L PRC: z = 0.12), his PHC was the only MTL structure that showed bilateral volume reduction (R and L PHC: z < −1.2). The latter might be the cause of his deficits in familiarity (rather than the unilateral PRC lesion that initially brought him to our attention). This again reinforces the need for larger patient cohorts in order to have sufficient variability in the laterality of MTL damage.

Finally, the present study did not examine other brain abnormalities that may be associated with MTL damage, such as lesion-induced changes in functional activity or connectivity. Thus, whether the effects of MTL damage on recollection and familiarity that we noted here are better explained by abnormalities in broader functional networks involving MTL regions (Argyropoulos et al., 2019) needs further investigation. Moreover, our scanning protocols did not allow us to investigate the integrity of smaller structures within the Papez circuit, which have also been implicated in memory, such as the thalamic nuclei and mammillary bodies (Aggleton and Brown, 1999; Kafkas et al., 2020; Tsivilis et al., 2008).

### Conclusion

We believe that our data provide the most compelling support yet for dual-process models of recognition memory, in which recollection and familiarity depend on different MTL structures. By capitalizing on the variability of damage across patients with MTL pathology, our study overcomes the limitations of single-case approaches, and lends new emphasis to the PHC as necessary for familiarity. Future studies of even larger patient groups, ideally across centers and using multiple, common paradigms and material-types, will hopefully further dissect the contributions of different MTL regions to memory.

## Data availability

Scripts for statistical analysis, behavioral and volumetric data are publicly available at: https://osf.io/a82ht

## Acknowledgments

We are very grateful to the participants who took part in this study. This research was supported by a Medical Research Council Clinician Scientist Fellowship to CRB (MR/K010395/1). R.N.H. is supported by UK Medical Research Council programme grant SUAG/046 G101400.Author information

## Author information

### Affiliations

*Memory Research Group, Nuffield Department of Clinical Neurosciences, University of Oxford, Oxford, UK; Level 6, West Wing, John Radcliffe Hospital, Oxford, OX3 9DU, UK*.

Georgios P. D. Argyropoulos, Carola Dell’Acqua, Emily Butler, Clare Loane, Adriana Roca-Fernandez, Azhaar Almozel, Nikolas Drummond, Carmen Lage-Martinez, Christopher R. Butler

*University of Stirling, Division of Psychology, Faculty of Natural Sciences, Stirling, UK*

Georgios P. D. Argyropoulos

*Department of General Psychology & Padova Neuroscience Center, University of Padova, Padova, Italy*

Carola Dell’Acqua

*Maurice Wohl Clinical Neuroscience Institute, Basic and Clinical Neuroscience*

*Department, King’s College London, London, UK*

Clare Loane

*Cardiff University, School of Biosciences, Cardiff, UK*

Azhaar Almozel

*Department of Zoology, University of Cambridge, Cambridge, UK*

Nikolas Drummond

*Valdecilla Biomedical Research Institute, University Hospital Marqués de Valdecilla*.

*Carmen Lage-Martinez*

*MRC Cognition & Brain Sciences Unit, and Department of Psychiatry, Cambridge, UK*

Elisa Cooper, Richard N. Henson

*Department of Brain Sciences, Imperial College London, London, UK*

Christopher R. Butler

*Departamento de Neurología, Pontificia Universidad Católica de Chile, Santiago, Chile*.

Christopher R. Butler

## Contributions

CRB conceptualized the study.

EB, ND, and CRB designed the experiments.

GPDA, CDA, EB, CL, ARF, AA, ND, CLM, and CRB collected the data.

GPDA, RNH, CDA, EB, CL, ARF, AA, and EC analyzed the data.

GPDA, RNH and CRB wrote the paper.

CRB provided clinical assessment.

CRB and RNH supervised the study.

## Corresponding author

Correspondence to Georgios P. D. Argyropoulos.

