## Supplementary Material for "Functional specialization of the medial temporal lobes in human recognition memory: dissociating effects of hippocampal vs parahippocampal damage"

### Supplementary Information

#### Supplementary table 1

Patients’ neuropsychological profile. Scores are age-scaled standardized scores (z), except for those marked with * (not age-scaled: based on the published means and standard deviations); **key:** **C&CT:** Camel and Cactus Test (Bozeat et al., 2000); **DC:** dot counting; **DKEFS:** Delis-Kaplan executive function system(Delis et al., 2001); **D&P:** Doors and People test (Baddeley et al., 1994); **DS:** digit span (WMS-III); **GNT:** Graded Naming Test (Mckenna and Warrington, 1980); **LNS:** letter-number switching (DKEFS); **MR:** Matrix Reasoning (WASI-II); **NART:** national adult reading test (Nelson and Willison, 1991); **PD:** position discrimination (VOSP); **p-FSIQ:** premorbid full-scale intelligence quotient (NART); **RMT:** Recognition Memory Test and Camden Memory Test for scenes (Warrington, 1984, 1996); **ROCFT:** Rey-Osterrieth Complex Figures Test (Rey, 1959); **SI:** Similarities (WASI-II); **VOSP:** Visual Object and Space Perception battery(Warrington and James, 1991); **WASI-II:** Wechsler Abbreviated Scale of Intelligence 2^nd^ Edition (Wechsler, 2011); **WL1:** word-list immediate recall (WMS-III); **WL2:** word-list delayed recall (WMS-III); **WL rec:** Word-list recognition test; **VC:** Vocabulary (WASI-II); **WMS-III:** Wechsler Memory Scale 3d edition (Wechsler, 1997); highlighted: z ≤ -1.67, i.e. the conventional cut-off point that corresponds to the 5^th^ %ile commonly employed to define impairment in neuropsychological assessment; **n/a:** not available (tests not administered due to scheduling conflicts).

| Test | NART | WASI-II | | | GNT * | C&CT * | VOSP | | | DKEFS | WMS-III | | | | | | RMT | | | ROCFT | | D&P | | | |
| --- | --- | --- | --- | --- | --- | --- | --- | --- | --- | --- | --- | --- | --- | --- | --- | --- | --- | --- | --- | --- | --- | --- | --- | --- | --- |
|  | p-FSIQ | MR | SI | VC |  |  | DC | CA | PD | LNS | DS | LM1 | LM2 | WL1 | WL2 | WL rec | Scenes | Faces | Words | Imm. | Del. | Names | People | Shapes | Doors |
| Function | Intelligence, Semantic Memory and Language | | | | | | Visuospatial | | | Executive | | Episodic Memory | | | | | | | | | | | | | |
| Patient |  | | | | | | | | | | | | | | | | | | | | | | | | |
| MH - PRC lesion | 0.80 | -0.75 | 0.67 | 0.88 | 0.39 | 0.02 | > 5^th^ %ile | | | -0.33 | 0.00 | n/a | | -0.67 | -0.33 | 1.00 | -1.12 | -2.33 | 0.00 | -0.75 | -0.82 | 2.67 | -0.33 | 0.33 | 0.67 |
| H1 -HPC lesion | 0.00 | -0.30 | -0.60 | 0.00 | -0.67 | 0.02 |  |  |  | -1.00 | -1.33 | -0.67 | 0.00 | -0.33 | 1.33 | 1.00 | -0.49 | -1.00 | 1.00 | -0.42 | 0.00 | 1.00 | -2.00 | 1.33 | 0.33 |
| H2 - HPC lesion | 1.20 | 0.30 | 1.00 | 0.60 | -0.10 | 0.99 |  |  |  | 0.00 | 0.33 | 1.00 | -0.67 | -0.67 | 0.33 | 1.00 | 1.68 | -0.67 | 0.67 | 0.40 | 0.00 | -1.67 | 0.67 | 1.33 | 2.33 |
| H3 - HPC lesion | 1.07 | 1.20 | 1.00 | 2.00 | -0.10 | 0.34 |  |  |  | -0.33 | 0.33 | -1.33 | -0.33 | 1.33 | -0.33 | -0.67 | 1.13 | 0.00 | -0.33 | 0.79 | 1.26 | -1.33 | -2.00 | -0.67 | 0.33 |
| H4 - HPC lesion | -0.13 | -1.20 | 0.70 | -0.80 | -0.83 | 0.67 |  |  |  | 1.33 | 0.33 | -1.00 | -2.33 | -2.00 | 0.00 | -0.33 | 0.09 | -0.33 | 0.00 | -3.00 | -3.00 | -2.33 | -0.33 | -1.00 | -1.00 |
| H5 - HPC lesion | 1.87 | 1.00 | 0.70 | 2.10 | 1.12 | 0.67 |  |  |  | 1.67 | 2.67 | 0.33 | -1.00 | -0.33 | 0.67 | -1.33 | n/a | 1.67 | 1.00 | 1.70 | 1.44 | 2.00 | -0.67 | 0.67 | 0.67 |
| H6 - HPC lesion | 1.53 | 0.10 | 1.80 | 3.00 | 0.63 | -0.64 |  |  |  | 0.33 | -0.33 | -2.00 | -2.67 | 0.00 | -1.33 | -1.33 | -1.84 | 2.33 | -1.33 | -2.16 | -3.00 | 0.67 | -1.67 | -2.00 | -1.67 |
| H7 - HPC lesion | 1.47 | -0.60 | 0.60 | 0.10 | -0.34 | 0.34 |  |  |  | 0.00 | -0.33 | -0.33 | -2.00 | 0.00 | 1.33 | 1.33 | -1.84 | 0.00 | 0.67 | -1.20 | -1.31 | -2.67 | -2.00 | -2.33 | -0.67 |
| H8 - HPC and PRC lesion | 1.13 | 1.70 | 1.50 | -1.00 | 0.88 | 0.99 |  |  |  | 1.33 | 2.00 | -1.67 | -1.67 | -2.00 | -1.33 | -2.00 | -0.83 | -2.33 | -1.33 | -3.00 | -1.51 | 0.00 | -1.67 | -1.33 | -0.67 |

#### Supplementary figure 1

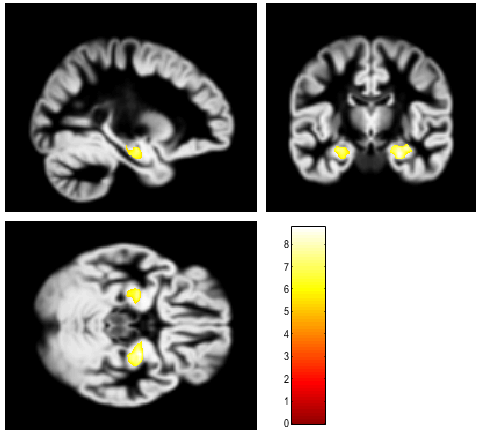

VBM results for contrast ‘CTRs (n=67) > HPC patients (n=8)’ on modulated GM tissue maps (reflecting GM volume). Volume reduction is noted bilaterally in the anterior HPC: Cluster 1: Right anterior HPC: cluster size: kE = 1865 voxels; peak coordinates: x=25, y=-11, z=-19 mm; t = 8.74; p-FWE < 0.0005; Cluster 2: Left anterior HPC: cluster size: kE = 863 voxels; peak coordinates: x=-25; y=-13; z=-19; t = 7.29 mm; p-FWE < 0.0005. Rest of ps: p-FWE > 0.15. Second-level between-subjects covariates: age at research scan; sex; study [see (Argyropoulos et al., 2019) for details]; total intracranial volume]. The voxels presented survive peak-level whole-brain FWE-correction (p < 0.05). Clusters are overlaid on an ICBM152 GM template (MNI space). Color-bar indicates t-values.

#### Supplementary figure 2

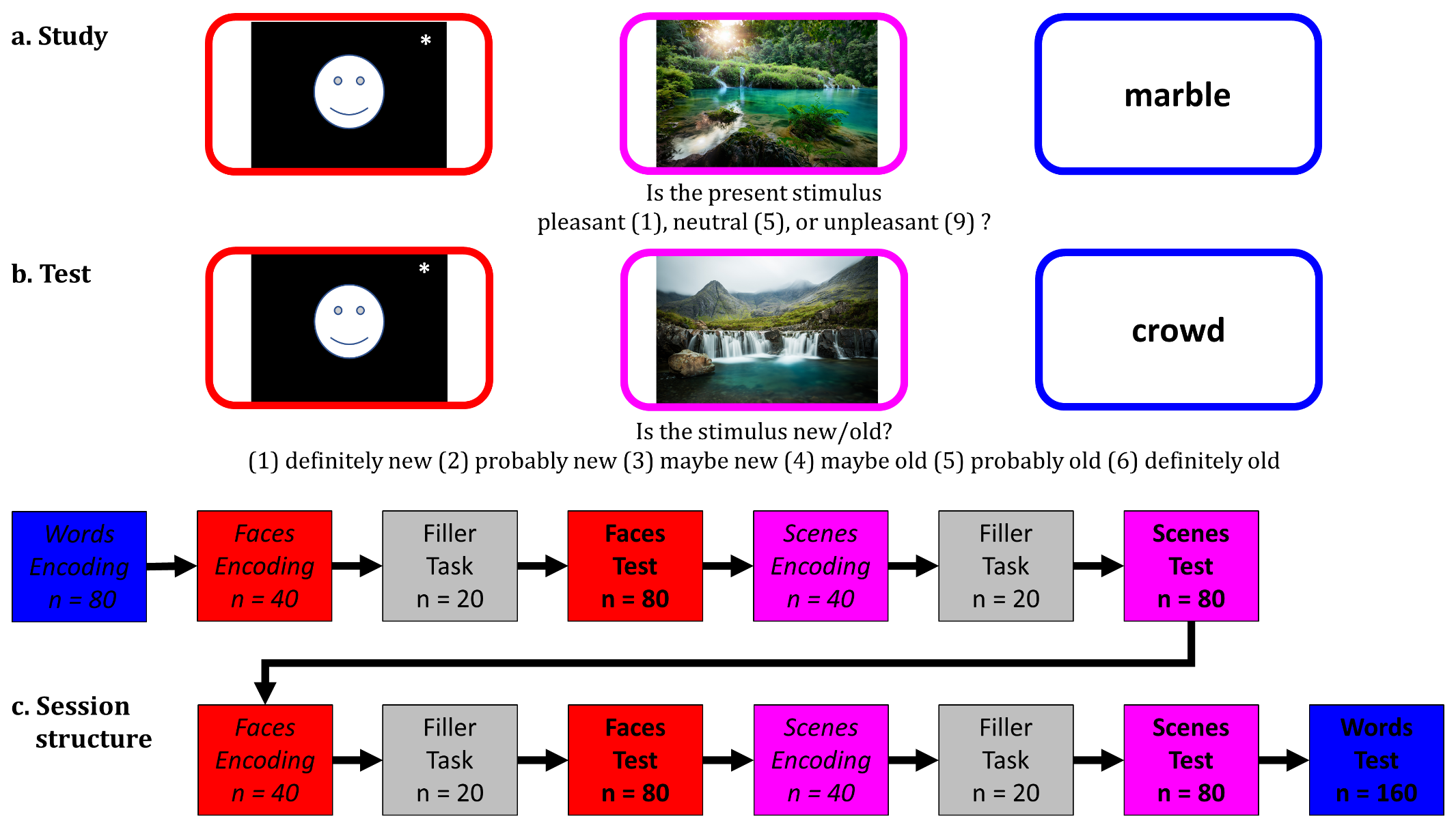

Paradigm 1 (ROC): Paradigm involving analysis of Receiver Operating Characteristics; a. Study phase: participants judged whether each stimulus was ‘pleasant’, ‘neutral’, or ‘unpleasant’; b. Test phase: participants were presented with each stimulus and were asked to judge, in a self-paced fashion, whether they have each stimulus before, rating their confidence on a scale from 1 to 6; c: session structure: the order of blocks was held constant across participants; the order of trials within each block was randomized for each session. Faces and Scenes were tested in two study-test blocks (separated by a filler task), whereas words were tested in a single pair of blocks at start and end of experiment, because memory for words is generally superior overall. blue: words; red: faces; green: scenes; grey: filler task; n: number of trials per block. *: **bioRχiv** **policy is to avoid the inclusion of photographs and any other identifying information of people, as verification of their consent is incompatible with the rapid and automated nature of preprint posting.**

#### Supplementary figure 3

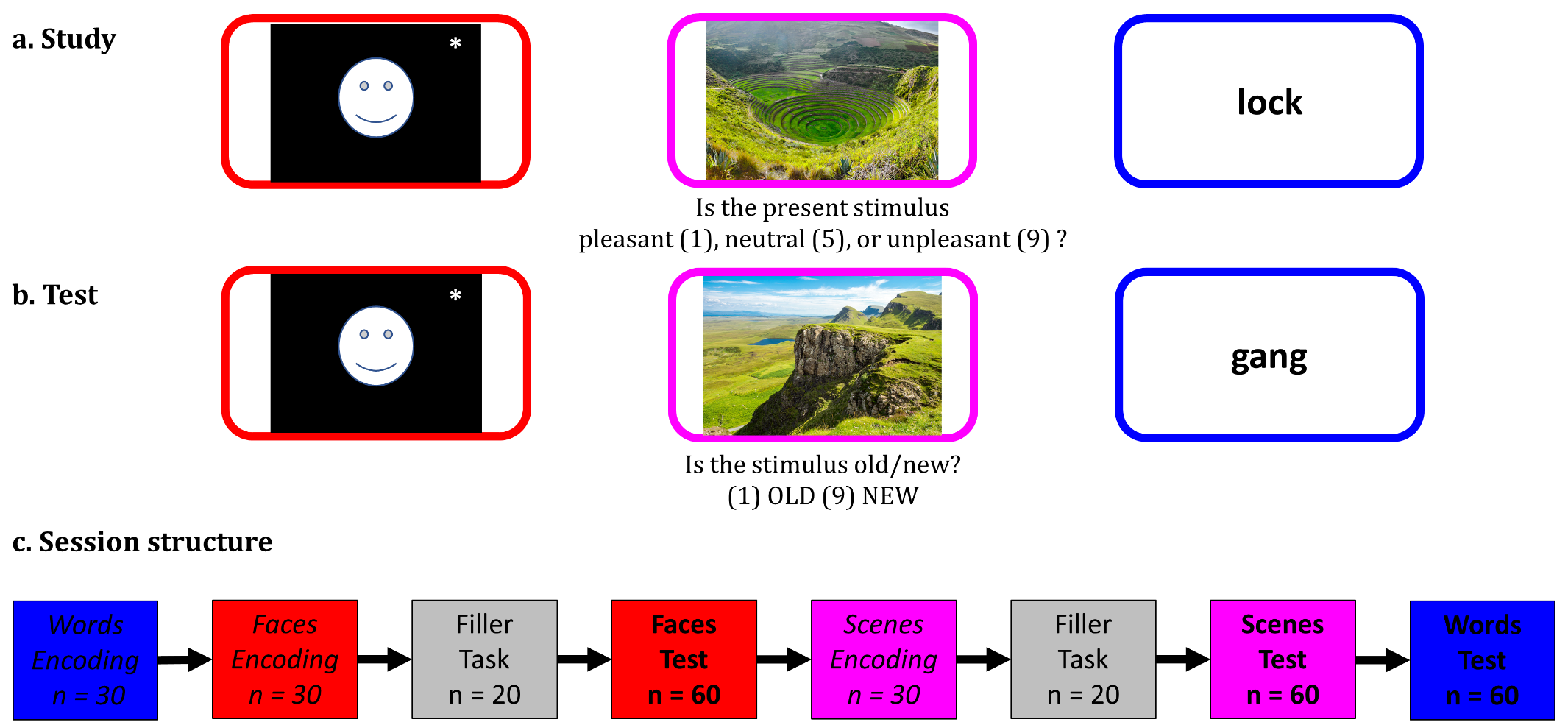

Paradigm 2 (RDP): Response Deadline Paradigm; a. Study phase: participants judged whether each stimulus was ‘pleasant’, ‘neutral’, or ‘unpleasant’; b. Test phase: participants were presented with each stimulus and were asked to judge whether they had encountered each stimulus before; c: session structure: the order of blocks was held constant across participants; the order of trials within each block was randomized for each session – see Supplementary Figure 2 legend for more details. The short versus long response deadline was manipulated across two sessions on different days (see Methods). blue: words; red: faces; green: scenes; grey: filler task; *n:* number of trials per block. *: **bioRχiv** **policy is to avoid the inclusion of photographs and any other identifying information of people, as verification of their consent is incompatible with the rapid and automated nature of preprint posting.**

#### Supplementary figure 4

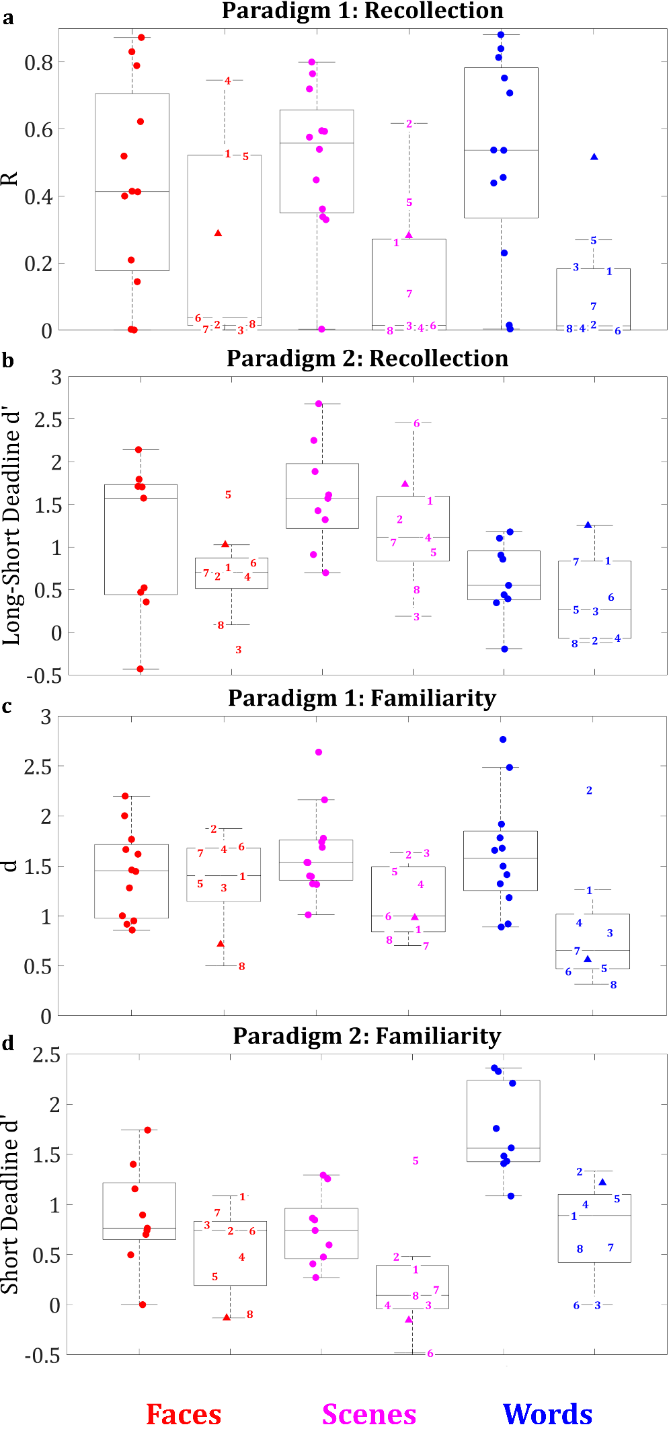

Recollection and familiarity estimates for CTRs and patients for face, scene, and word recognition memory in Paradigms 1 and 2; **a-b: recollection estimates; c-d: familiarity estimates;** in Paradigm 2 (RDP), familiarity is reflected by the sensitivity indices (d’) for the short deadline, whereas recollection by the difference between the d’ for the long response deadline and that for the short response deadline. Line in boxplots=median; bottom of box=25^th^ %ile; top=75^th^ %ile; whiskers: 1.5 * interquartile range; **key:** L, R: left/right hemisphere; ● CTRs : healthy controls; ▲: MH (patient with right PRC lesion); H1-7: patients with HPC but no PRC lesion; H8: patient with both HPC and right PRC lesions; **PRC:** perirhinal cortex; **HPC:** hippocampus;

#### Supplementary table 2

| Analysis | Main Effect / Interaction | df1 | df2 | MICE IMPUTATION | | | | | | | | | |
| --- | --- | --- | --- | --- | --- | --- | --- | --- | --- | --- | --- | --- | --- |
|  |  |  |  | 1 | | 2 | | 3 | | 4 | | 5 | |
|  |  |  |  | F | p | F | p | F | p | F | p | F | p |
| MH vs CTRs  (4-way ANOVA) | Group | 1 | 13 | 7.14 | 0.019 | 6.53 | 0.024 | 6.03 | 0.029 | 6.18 | 0.027 | 6.27 | 0.026 |
|  | Material-Type | 2 | 26 | 5.42 | 0.011 | 4.84 | 0.016 | 3.44 | 0.047 | 5.11 | 0.014 | 4.67 | 0.019 |
|  | Group*Material-Type | 2 | 26 | 0.69 | 0.512 | 0.20 | 0.818 | 0.42 | 0.663 | 0.47 | 0.629 | 0.45 | 0.640 |
|  | Process | 1 | 13 | 30.54 | < 0.001 | 22.13 | < 0.001 | 28.84 | < 0.001 | 28.63 | < 0.001 | 32.57 | < 0.001 |
|  | Group*Process | 1 | 13 | 5.46 | 0.036 | 4.05 | 0.065 | 5.70 | 0.033 | 5.35 | 0.038 | 5.92 | 0.030 |
|  | Paradigm | 1 | 13 | 5.89 | 0.031 | 6.45 | 0.025 | 5.57 | 0.035 | 0.97 | 0.343 | 10.54 | 0.006 |
|  | Group*Paradigm | 1 | 13 | 0.22 | 0.646 | 0.21 | 0.657 | 0.29 | 0.603 | 0.29 | 0.602 | 0.15 | 0.701 |
|  | Material-Type*Process | 2 | 26 | 16.10 | < 0.001 | 19.80 | < 0.001 | 14.61 | < 0.001 | 18.39 | < 0.001 | 14.00 | < 0.001 |
|  | Group*Material-Type*Process | 2 | 26 | 0.21 | 0.809 | 0.34 | 0.713 | 0.14 | 0.872 | 0.27 | 0.763 | 0.17 | 0.845 |
|  | Paradigm*Material-Type | 2 | 26 | 0.04 | 0.958 | 0.04 | 0.958 | 0.11 | 0.892 | 0.08 | 0.922 | 1.00 | 0.381 |
|  | Group*Paradigm*Material-Type | 2 | 26 | 1.65 | 0.212 | 0.97 | 0.393 | 1.28 | 0.295 | 1.81 | 0.184 | 1.08 | 0.354 |
|  | Paradigm*Process | 1 | 13 | 28.79 | < 0.001 | 57.11 | < 0.001 | 29.27 | < 0.001 | 34.01 | < 0.001 | 28.30 | < 0.001 |
|  | Group*Paradigm*Process | 1 | 13 | 0.48 | 0.503 | 0.32 | 0.582 | 0.27 | 0.612 | 0.36 | 0.559 | 0.48 | 0.501 |
|  | Paradigm*Material-Type*Process | 2 | 26 | 13.31 | < 0.001 | 20.99 | < 0.001 | 15.23 | < 0.001 | 12.37 | < 0.001 | 16.75 | < 0.001 |
|  | Group*Paradigm*Material-Type*Process | 2 | 26 | 0.08 | 0.924 | 0.06 | 0.945 | 0.10 | 0.906 | 0.16 | 0.852 | 0.06 | 0.941 |
| MH vs CTRs: Recollection  (3-way ANOVA) | Group | 1 | 13 | 0.04 | 0.837 | 0.01 | 0.911 | 0.01 | 0.915 | 0.05 | 0.828 | 0.02 | 0.884 |
|  | Material-Type | 2 | 26 | 6.76 | 0.004 | 6.18 | 0.007 | 5.66 | 0.009 | 8.21 | 0.002 | 6.81 | 0.004 |
|  | Group*Material-Type | 2 | 26 | 0.53 | 0.597 | 0.40 | 0.677 | 0.41 | 0.671 | 0.77 | 0.474 | 0.46 | 0.637 |
|  | Paradigm | 1 | 13 | 29.78 | < 0.001 | 42.54 | < 0.001 | 32.30 | < 0.001 | 19.52 | < 0.001 | 33.09 | < 0.001 |
|  | Group*Paradigm | 1 | 13 | 0.62 | 0.445 | 0.41 | 0.533 | 0.50 | 0.492 | 0.55 | 0.472 | 0.54 | 0.477 |
|  | Paradigm*Material-Type | 2 | 26 | 13.69 | < 0.001 | 14.92 | < 0.001 | 11.19 | < 0.001 | 9.69 | < 0.001 | 10.03 | < 0.001 |
|  | Group*Paradigm*Material-Type | 2 | 26 | 0.43 | 0.655 | 0.39 | 0.682 | 0.25 | 0.785 | 0.27 | 0.765 | 0.22 | 0.803 |
| MH vs CTRs: Familiarity  (3-way ANOVA) | Group | 1 | 13 | 13.74 | 0.003 | 10.00 | 0.008 | 14.31 | 0.002 | 11.06 | 0.005 | 12.16 | 0.004 |
|  | Material-Type | 2 | 26 | 24.26 | < 0.001 | 28.17 | < 0.001 | 16.09 | < 0.001 | 18.05 | < 0.001 | 16.03 | < 0.001 |
|  | Group*Material-Type | 2 | 26 | 0.03 | 0.972 | 0.07 | 0.933 | 0.04 | 0.959 | 0.010 | 0.994 | 0.03 | 0.974 |
|  | Paradigm | 1 | 13 | 9.89 | 0.008 | 23.06 | < 0.001 | 11.46 | 0.005 | 13.75 | 0.003 | 9.21 | 0.010 |
|  | Group*Paradigm | 1 | 13 | 0.09 | 0.765 | 0.02 | 0.879 | 0.02 | 0.879 | < 0.01 | 0.967 | 0.17 | 0.687 |
|  | Paradigm*Material-Type | 2 | 26 | 8.16 | 0.002 | 10.04 | < 0.001 | 9.65 | < 0.001 | 9.02 | 0.001 | 14.04 | < 0.001 |
|  | Group*Paradigm*Material-Type | 2 | 26 | 0.48 | 0.624 | 0.49 | 0.619 | 0.67 | 0.519 | 0.81 | 0.455 | 0.45 | 0.640 |

ANOVAs (between-participants independent variables: Group (MH (PRC lesion) vs. CTRs); within-participants independent variables: Paradigm (ROC vs. RDP); Process (Familiarity vs. Recollection); Material-Type (Faces, Scenes, Words)) on (recollection and familiarity) estimates for MH (focal PRC lesion) and CTRs for faces, scenes, and words, in the ROC and the RDP paradigms. These missing values were imputed using “Multiple Imputation with Chained Equations” implemented in the R function “mice”. Five imputations were created (MICE 1-5). shaded cells: significant (p < 0.05) main effects and interactions of interest.

#### Supplementary table 3

The ANOVA for the HPC group versus CTRs did not show a Group*Process interaction. Nevertheless, we observed a significant Group*Material-Type interaction (F(2,38) = 9.83, p < 0.001), along with a borderline three-way interaction between Group, Process and Material-Type (F(2, 38) =3.02, p = 0.061). The follow-up ANOVA for Recollection showed only a main effect of Group (F(1, 19) = 7.89, p = 0.011; Group*Material-Type: F(2,38) = 0.02, p = 0.981), providing no evidence that recollection in HPC patients is differentially impaired across Material-Types. However, the follow-up ANOVA for Familiarity also showed a significant interaction between Group and Material-Type (F(2,38) = 10.44, p < 0.001; Group: F(1,19) = 20.14, p < 0.001). This was further decomposed by examining Familiarity for each Material-Type separately. Faces showed no significant effects (F < 0.5), suggesting no impairment of this group on Face Familiarity. However, ANOVAs on both Scene and Word Familiarity showed a main effect of Group, with patients scoring lower than CTRs (F(1,19) = 12.58, p = 0.002 and F(1,19) = 24.18, p < 0.001, respectively).

| Analysis | Main Effect / Interaction | df1 | df2 | MICE IMPUTATION | | | | | | | | | |
| --- | --- | --- | --- | --- | --- | --- | --- | --- | --- | --- | --- | --- | --- |
|  |  |  |  | 1 | | 2 | | 3 | | 4 | | 5 | |
|  |  |  |  | F | P | F | p | F | P | F | p | F | p |
| HPC (H1-H7) vs. CTRs  (4-way ANOVA) | Group | 1 | 19 | 31.42 | < 0.001 | 30.45 | < 0.001 | 27.76 | < 0.001 | 27.86 | < 0.001 | 29.52 | < 0.001 |
|  | Material-Type | 2 | 38 | 0.81 | 0.453 | 0.91 | 0.410 | 0.50 | 0.611 | 0.57 | 0.571 | 0.77 | 0.470 |
|  | Group*Material-Type | 2 | 38 | 9.83 | < 0.001 | 7.81 | 0.001 | 7.12 | 0.002 | 9.28 | 0.001 | 8.49 | 0.001 |
|  | Process | 1 | 19 | 47.46 | < 0.001 | 37.13 | < 0.001 | 45.93 | < 0.001 | 45.45 | < 0.001 | 49.85 | < 0.001 |
|  | Group*Process | 1 | 19 | 1.09 | 0.310 | 0.72 | 0.408 | 0.45 | 0.513 | 0.74 | 0.399 | 1.01 | 0.327 |
|  | Paradigm | 1 | 19 | 1.93 | 0.181 | 2.15 | 0.159 | 1.67 | 0.212 | 0.19 | 0.667 | 3.54 | 0.076 |
|  | Group*Paradigm | 1 | 19 | 2.84 | 0.109 | 3.04 | 0.098 | 2.67 | 0.119 | 0.75 | 0.398 | 4.34 | 0.051 |
|  | Material-Type*Process | 2 | 38 | 14.10 | < 0.001 | 17.15 | < 0.001 | 13.03 | < 0.001 | 14.68 | < 0.001 | 13.13 | < 0.001 |
|  | Group*Material-Type*Process | 2 | 38 | 3.02 | 0.061 | 3.64 | 0.036 | 2.40 | 0.105 | 3.76 | 0.032 | 2.49 | 0.096 |
|  | Paradigm*Material-Type | 2 | 38 | 0.28 | 0.759 | 0.39 | 0.680 | 0.61 | 0.551 | 0.45 | 0.641 | 0.97 | 0.389 |
|  | Group*Paradigm*Material-Type | 2 | 38 | 1.10 | 0.344 | 0.51 | 0.604 | 0.58 | 0.563 | 1.04 | 0.363 | 0.92 | 0.406 |
|  | Paradigm*Process | 1 | 19 | 57.55 | < 0.001 | 95.79 | < 0.001 | 56.66 | < 0.001 | 64.61 | < 0.001 | 56.80 | < 0.001 |
|  | Group*Paradigm*Process | 1 | 19 | 1.43 | 0.247 | 0.56 | 0.463 | 0.68 | 0.420 | 0.93 | 0.347 | 1.45 | 0.244 |
|  | Paradigm*Material-Type*Process | 2 | 38 | 17.26 | < 0.001 | 24.66 | < 0.001 | 19.24 | < 0.001 | 16.32 | < 0.001 | 20.78 | < 0.001 |
|  | Group*Paradigm*Material-Type*Process | 2 | 38 | 0.44 | 0.650 | 0.79 | 0.462 | 0.34 | 0.715 | 0.18 | 0.839 | 0.53 | 0.593 |
| HPC (H1-H7) vs. CTRs: Recollection  (3-way ANOVA) | Group | 1 | 19 | 7.89 | 0.011 | 8.23 | 0.010 | 9.36 | 0.006 | 8.81 | 0.008 | 9.49 | 0.006 |
|  | Material-Type | 2 | 38 | 11.65 | < 0.001 | 10.63 | < 0.001 | 10.29 | < 0.001 | 13.43 | < 0.001 | 11.61 | < 0.001 |
|  | Group*Material-Type | 2 | 38 | 0.02 | 0.981 | 0.03 | 0.976 | 0.11 | 0.899 | 0.13 | 0.881 | 0.04 | 0.957 |
|  | Paradigm | 1 | 19 | 40.58 | < 0.001 | 53.12 | < 0.001 | 43.17 | < 0.001 | 29.36 | < 0.001 | 43.95 | 0.000 |
|  | Group*Paradigm | 1 | 19 | 0.02 | 0.883 | 0.37 | 0.551 | 0.10 | 0.751 | 0.00 | 0.978 | 0.09 | 0.768 |
|  | Paradigm*Material-Type | 2 | 38 | 17.78 | < 0.001 | 19.25 | < 0.001 | 15.61 | < 0.001 | 13.94 | < 0.001 | 14.41 | < 0.001 |
|  | Group*Paradigm*Material-Type | 2 | 38 | 1.13 | 0.334 | 1.08 | 0.351 | 0.57 | 0.568 | 0.62 | 0.541 | 0.47 | 0.631 |
| HPC (H1-H7) vs. CTRs: Familiarity  (3-way ANOVA) | Group | 1 | 19 | 20.14 | < 0.001 | 16.49 | 0.001 | 18.18 | < 0.001 | 16.04 | 0.001 | 18.84 | < 0.001 |
|  | Material-Type | 2 | 38 | 9.62 | < 0.001 | 12.61 | < 0.001 | 7.33 | 0.002 | 8.20 | 0.001 | 7.78 | 0.001 |
|  | Group*Material-Type | 2 | 38 | 10.44 | < 0.001 | 12.07 | < 0.001 | 8.01 | 0.001 | 9.37 | < 0.001 | 8.16 | 0.001 |
|  | Paradigm | 1 | 19 | 27.59 | < 0.001 | 46.74 | < 0.001 | 28.95 | < 0.001 | 31.82 | < 0.001 | 27.33 | < 0.001 |
|  | Group*Paradigm | 1 | 19 | 4.49 | 0.048 | 3.94 | 0.062 | 2.89 | 0.106 | 1.97 | 0.177 | 5.83 | 0.026 |
|  | Paradigm*Material-Type | 2 | 38 | 10.27 | < 0.001 | 12.24 | < 0.001 | 11.80 | < 0.001 | 11.14 | < 0.001 | 16.19 | < 0.001 |
|  | Group*Paradigm*Material-Type | 2 | 38 | 0.27 | 0.769 | 0.39 | 0.682 | 0.27 | 0.765 | 0.18 | 0.834 | 0.79 | 0.464 |
| HPC (H1-H7) vs. CTRs: Faces  (3-way ANOVA) | Group | 1 | 19 | 4.35 | 0.051 | 2.16 | 0.158 | 2.40 | 0.138 | 3.52 | 0.076 | 4.44 | 0.049 |
|  | Process | 1 | 19 | 11.91 | 0.003 | 11.14 | 0.003 | 12.27 | 0.002 | 12.48 | 0.002 | 12.68 | 0.002 |
|  | Group*Process | 1 | 19 | 0.70 | 0.413 | 0.89 | 0.359 | 0.91 | 0.353 | 1.65 | 0.214 | 0.59 | 0.452 |
|  | Paradigm | 1 | 19 | 0.59 | 0.453 | 0.21 | 0.656 | 0.06 | 0.808 | 0.02 | 0.902 | 0.27 | 0.610 |
|  | Group*Paradigm | 1 | 19 | 7.98 | 0.011 | 4.07 | 0.058 | 3.96 | 0.061 | 2.41 | 0.137 | 5.41 | 0.031 |
|  | Paradigm*Process | 1 | 19 | 20.55 | < 0.001 | 34.27 | < 0.001 | 26.20 | < 0.001 | 21.61 | < 0.001 | 24.00 | < 0.001 |
|  | Group*Paradigm*Process | 1 | 19 | 0.05 | 0.827 | 0.00 | 0.978 | 0.07 | 0.801 | 0.06 | 0.817 | 0.17 | 0.688 |
| HPC (H1-H7) vs. CTRs: Scenes  (3-way ANOVA) | Group | 1 | 19 | 19.01 | < 0.001 | 18.88 | < 0.001 | 16.86 | 0.001 | 19.91 | < 0.001 | 16.30 | 0.001 |
|  | Process | 1 | 19 | 2.20 | 0.154 | 0.87 | 0.364 | 1.43 | 0.246 | 2.34 | 0.142 | 1.76 | 0.200 |
|  | Group*Process | 1 | 19 | 0.71 | 0.412 | 0.20 | 0.660 | 0.38 | 0.543 | 0.77 | 0.391 | 0.52 | 0.480 |
|  | Paradigm | 1 | 19 | 1.48 | 0.238 | 1.71 | 0.207 | 1.87 | 0.188 | 0.44 | 0.513 | 1.70 | 0.208 |
|  | Group*Paradigm | 1 | 19 | 0.27 | 0.612 | 0.38 | 0.546 | 0.38 | 0.545 | 0.03 | 0.876 | 0.30 | 0.591 |
|  | Paradigm*Process | 1 | 19 | 94.34 | < 0.001 | 114.04 | < 0.001 | 92.34 | < 0.001 | 111.51 | < 0.001 | 78.52 | < 0.001 |
|  | Group*Paradigm*Process | 1 | 19 | 0.33 | 0.575 | 0.01 | 0.946 | 0.06 | 0.816 | 0.37 | 0.552 | 0.10 | 0.760 |
| HPC (H1-H7) vs. CTRs: Words  (3-way ANOVA) | Group | 1 | 19 | 44.46 | < 0.001 | 41.97 | < 0.001 | 47.13 | < 0.001 | 31.13 | < 0.001 | 37.81 | < 0.001 |
|  | Process | 1 | 19 | 75.82 | < 0.001 | 58.33 | < 0.001 | 66.32 | < 0.001 | 65.54 | < 0.001 | 65.97 | < 0.001 |
|  | Group*Process | 1 | 19 | 5.73 | 0.027 | 4.99 | 0.038 | 3.60 | 0.073 | 4.62 | 0.045 | 4.70 | 0.043 |
|  | Paradigm | 1 | 19 | 1.24 | 0.279 | 1.42 | 0.248 | 1.27 | 0.275 | 0.27 | 0.610 | 5.34 | 0.032 |
|  | Group*Paradigm | 1 | 19 | 1.00 | 0.331 | 1.15 | 0.298 | 1.05 | 0.318 | 0.33 | 0.572 | 3.48 | 0.078 |
|  | Paradigm*Process | 1 | 19 | 0.15 | 0.705 | 0.66 | 0.426 | 0.65 | 0.431 | 1.23 | 0.281 | 0.11 | 0.742 |
|  | Group*Paradigm*Process | 1 | 19 | 2.53 | 0.128 | 2.77 | 0.113 | 1.34 | 0.262 | 1.30 | 0.269 | 3.01 | 0.099 |
| HPC (H1-H7) vs. CTRs: Familiarity: Faces  (2-way ANOVA) | Group | 1 | 19 | 0.26 | 0.616 | 0.07 | 0.795 | 0.10 | 0.751 | 0.00 | 0.976 | 0.35 | 0.560 |
|  | Paradigm | 1 | 19 | 18.24 | < 0.001 | 24.80 | < 0.001 | 26.63 | < 0.001 | 23.34 | < 0.001 | 18.74 | < 0.001 |
|  | Group*Paradigm | 1 | 19 | 2.27 | 0.148 | 1.64 | 0.216 | 2.48 | 0.132 | 1.56 | 0.227 | 2.78 | 0.112 |
| HPC (H1-H7) vs. CTRs: Familiarity: Scenes  (2-way ANOVA) | Group | 1 | 19 | 12.58 | 0.002 | 12.35 | 0.002 | 12.30 | 0.002 | 13.45 | 0.002 | 11.08 | 0.004 |
|  | Paradigm | 1 | 19 | 37.66 | < 0.001 | 41.22 | < 0.001 | 34.53 | < 0.001 | 39.80 | < 0.001 | 45.06 | < 0.001 |
|  | Group*Paradigm | 1 | 19 | 0.58 | 0.457 | 0.16 | 0.690 | 0.30 | 0.591 | 0.25 | 0.620 | 0.39 | 0.540 |
| HPC (H1-H7) vs. CTRs: Familiarity: Words  (2-way ANOVA) | Group | 1 | 19 | 24.18 | < 0.001 | 19.48 | < 0.001 | 19.95 | < 0.001 | 16.57 | 0.001 | 19.54 | < 0.001 |
|  | Paradigm | 1 | 19 | 0.08 | 0.780 | 0.01 | 0.919 | 0.01 | 0.935 | 0.26 | 0.616 | 0.86 | 0.366 |
|  | Group*Paradigm | 1 | 19 | 2.33 | 0.144 | 2.94 | 0.103 | 1.86 | 0.189 | 1.26 | 0.275 | 4.78 | 0.042 |
| HPC (H1-H7) vs. CTRs: Recollection: Faces  (2-way ANOVA) | Group | 1 | 19 | 2.26 | 0.149 | 1.74 | 0.203 | 2.14 | 0.160 | 3.45 | 0.079 | 2.21 | 0.154 |
|  | Paradigm | 1 | 19 | 16.59 | 0.001 | 21.22 | < 0.001 | 14.84 | 0.001 | 11.27 | 0.003 | 17.35 | 0.001 |
|  | Group*Paradigm | 1 | 19 | 0.75 | 0.398 | 1.21 | 0.285 | 0.54 | 0.473 | 0.31 | 0.587 | 0.58 | 0.454 |
| HPC (H1-H7) vs. CTRs: Recollection: Scenes  (2-way ANOVA) | Group | 1 | 19 | 3.71 | 0.069 | 4.79 | 0.041 | 4.13 | 0.056 | 2.84 | 0.109 | 4.58 | 0.046 |
|  | Paradigm | 1 | 19 | 63.93 | < 0.001 | 67.10 | < 0.001 | 77.20 | < 0.001 | 59.66 | < 0.001 | 53.63 | < 0.001 |
|  | Group*Paradigm | 1 | 19 | 0.00 | 0.952 | 0.24 | 0.629 | 0.06 | 0.805 | 0.09 | 0.774 | 0.01 | 0.935 |
| HPC (H1-H7) vs. CTRs: Recollection: Words  (2-way ANOVA) | Group | 1 | 19 | 9.02 | 0.007 | 9.51 | 0.006 | 18.14 | < 0.001 | 13.51 | 0.002 | 11.85 | 0.003 |
|  | Paradigm | 1 | 19 | 1.90 | 0.184 | 2.79 | 0.112 | 2.16 | 0.158 | 1.91 | 0.183 | 3.68 | 0.070 |
|  | Group*Paradigm | 1 | 19 | 0.98 | 0.335 | 0.56 | 0.464 | 0.26 | 0.615 | 0.42 | 0.527 | 0.17 | 0.686 |

ANOVAs (between-participants independent variables: Group (HPC lesion (H1-H7), CTRs); within-participants independent variables: Paradigm (ROC, RDP); Process (familiarity, recollection); Material-Type (faces, scenes, words)) on (recollection and familiarity) estimates for MH (focal PRC lesion) and CTRs for faces, scenes, and words, in the ROC and the RDP paradigms. These missing values were imputed using “Multiple Imputation with Chained Equations” implemented in the R function “mice”. Five imputations were created (MICE 1-5). shaded cells: significant (p < 0.05) main effects and interactions of interest.

#### Supplementary table 4

| # Linear Model – Fixed Effects | **Effects/Interactions** | **df1** | **df2** | **F** | **P** |
| --- | --- | --- | --- | --- | --- |
| **1 – Group, Paradigm, Process, Material-Type** | Group | 1 | 6 | < 0.01 | 0.995 |
|  | Paradigm | 1 | 66 | 0.16 | 0.686 |
|  | Process | 1 | 66 | 0.42 | 0.519 |
|  | Material-Type | 2 | 66 | 10.51 | < 0.001 |
|  | Group*Paradigm | 1 | 66 | 0.73 | 0.397 |
|  | Group*Process | 1 | 66 | 8.52 | 0.005 |
|  | Paradigm*Process | 1 | 66 | 9.65 | 0.003 |
|  | Group*Material-Type | 2 | 66 | 3.20 | 0.047 |
|  | Paradigm*Material-Type | 2 | 66 | 0.45 | 0.638 |
|  | Process*Material-Type | 2 | 66 | 2.97 | 0.058 |
|  | Group*Paradigm*Process | 1 | 66 | < 0.01 | > 0.999 |
|  | Group*Paradigm*Material-Type | 2 | 66 | 0.55 | 0.581 |
|  | Group*Process*Material-Type | 2 | 66 | 0.09 | 0.914 |
|  | Paradigm*Process*Material-Type | 2 | 66 | 0.70 | 0.502 |
|  | Group*Paradigm*Process*Material-Type | 2 | 66 | 0.32 | 0.724 |
| 2 (Recollection) – Group, Paradigm | Group | 1 | 6 | 3.62 | 0.106 |
|  | Paradigm | 1 | 6 | 3.36 | 0.117 |
|  | Group*Paradigm | 1 | 6 | 0.33 | 0.584 |
| 3 (Familiarity) – Group, Paradigm | Group | 1 | 6 | 1.84 | 0.224 |
|  | Paradigm | 1 | 6 | 4.93 | 0.068 |
|  | Group*Paradigm | 1 | 6 | 0.29 | 0.609 |
| 4 (Faces) – Group, Paradigm | Group | 1 | 6 | 5.63 | 0.055 |
|  | Paradigm | 1 | 6 | 1.20 | 0.315 |
|  | Group*Paradigm | 1 | 6 | 0.18 | 0.689 |
| 5 (Scenes) – Group, Paradigm | Group | 1 | 6 | 0.05 | 0.838 |
|  | Paradigm | 1 | 6 | 0.35 | 0.577 |
|  | Group*Paradigm | 1 | 6 | 0.01 | 0.917 |
| 6 (Words) – Group, Paradigm | Group | 1 | 6 | 3.60 | 0.107 |
|  | Paradigm | 1 | 6 | 0.71 | 0.432 |
|  | Group*Paradigm | 1 | 6 | 3.70 | 0.103 |

We added both MH and the 7 HPC cases (H1-7) to a single linear model, which was fit to patients’ Z-scores relative to CTRs. The model included fixed effects of: Group (PRC vs HPC lesion); Paradigm (ROC vs RDP); Process (Recollection vs Familiarity) and Material-Type (Faces, Scenes, Words); shaded cells: significant (p < 0.05) main effects and interactions of interest.

#### Supplementary table 5

| **Model** | **1** | **df1** | **df2** | **F** | **p** |
| --- | --- | --- | --- | --- | --- |
| HPC | ROI Laterality | 1 | 167.71 | 0.02 | 0.889 |
|  | Volume | 1 | 28.53 | 0.89 | 0.354 |
|  | Paradigm | 1 | 160.33 | 4.59 | 0.034 |
|  | Process | 1 | 160.33 | 25.90 | < 0.001 |
|  | Material | 2 | 160.33 | 0.92 | 0.399 |
|  | ROI Laterality*Volume | 1 | 166.17 | 0.00 | 0.999 |
|  | ROI Laterality*Paradigm | 1 | 160.33 | 0.10 | 0.752 |
|  | Volume*Paradigm | 1 | 160.33 | 5.69 | 0.018 |
|  | ROI Laterality*Process | 1 | 160.33 | 0.37 | 0.541 |
|  | Volume*Process | 1 | 160.33 | 17.93 | < 0.001 |
|  | Paradigm*Process | 1 | 160.33 | 2.16 | 0.144 |
|  | ROI Laterality*Material | 2 | 160.33 | 0.10 | 0.903 |
|  | Volume*Material | 2 | 160.33 | 2.43 | 0.091 |
|  | Paradigm*Material | 2 | 160.33 | 1.75 | 0.178 |
|  | Process*Material | 2 | 160.33 | 0.49 | 0.611 |
|  | ROI Laterality*Volume*Paradigm | 1 | 160.33 | 0.02 | 0.894 |
|  | ROI Laterality*Volume*Process | 1 | 160.33 | 0.09 | 0.770 |
|  | ROI Laterality*Paradigm*Process | 1 | 160.33 | 0.29 | 0.592 |
|  | Volume*Paradigm*Process | 1 | 160.33 | 0.31 | 0.580 |
|  | ROI Laterality*Volume*Material | 2 | 160.33 | 0.23 | 0.793 |
|  | ROI Laterality*Paradigm*Material | 2 | 160.33 | 0.13 | 0.881 |
|  | Volume*Paradigm*Material | 2 | 160.33 | 1.53 | 0.220 |
|  | ROI Laterality*Process*Material | 2 | 160.33 | 0.71 | 0.493 |
|  | Volume*Process*Material | 2 | 160.33 | 0.23 | 0.797 |
|  | Paradigm*Process*Material | 2 | 160.33 | 0.07 | 0.932 |
|  | ROI Laterality*Volume*Paradigm*Process | 1 | 160.33 | 0.30 | 0.586 |
|  | ROI Laterality*Volume*Paradigm*Material | 2 | 160.33 | 0.10 | 0.907 |
|  | ROI Laterality*Volume*Process*Material | 2 | 160.33 | 0.86 | 0.425 |
|  | ROI Laterality*Paradigm*Process*Material | 2 | 160.33 | 0.05 | 0.949 |
|  | Volume*Paradigm*Process*Material | 2 | 160.33 | 0.08 | 0.927 |
|  | ROI Laterality*Volume*Paradigm*Process*Material | 2 | 160.33 | 0.07 | 0.930 |
| PRC | ROI Laterality | 1 | 167.23 | 0.01 | 0.935 |
|  | Volume | 1 | 133.63 | 0.01 | 0.906 |
|  | Paradigm | 1 | 160.4 | 0.69 | 0.407 |
|  | Process | 1 | 160.4 | 4.76 | 0.031 |
|  | Material | 2 | 160.4 | 11.57 | < 0.001 |
|  | ROI Laterality*Volume | 1 | 154.86 | 0.00 | 0.945 |
|  | ROI Laterality*Paradigm | 1 | 160.4 | 0.09 | 0.766 |
|  | Volume*Paradigm | 1 | 160.4 | 1.30 | 0.256 |
|  | ROI Laterality*Process | 1 | 160.4 | 1.20 | 0.276 |
|  | Volume*Process | 1 | 160.4 | 0.18 | 0.668 |
|  | Paradigm*Process | 1 | 160.4 | 14.14 | < 0.001 |
|  | ROI Laterality*Material | 2 | 160.4 | 0.98 | 0.377 |
|  | Volume*Material | 2 | 160.4 | 0.55 | 0.578 |
|  | Paradigm*Material | 2 | 160.4 | 0.51 | 0.601 |
|  | Process*Material | 2 | 160.4 | 2.40 | 0.094 |
|  | ROI Laterality*Volume*Paradigm | 1 | 160.4 | 0.36 | 0.552 |
|  | ROI Laterality*Volume*Process | 1 | 160.4 | 2.17 | 0.143 |
|  | ROI Laterality*Paradigm*Process | 1 | 160.4 | 0.02 | 0.883 |
|  | Volume*Paradigm*Process | 1 | 160.4 | 0.30 | 0.583 |
|  | ROI Laterality*Volume*Material | 2 | 160.4 | 1.64 | 0.197 |
|  | ROI Laterality*Paradigm*Material | 2 | 160.4 | 0.22 | 0.805 |
|  | Volume*Paradigm*Material | 2 | 160.4 | 0.07 | 0.933 |
|  | ROI Laterality*Process*Material | 2 | 160.4 | 0.23 | 0.793 |
|  | Volume*Process*Material | 2 | 160.4 | 0.17 | 0.846 |
|  | Paradigm*Process*Material | 2 | 160.4 | 2.60 | 0.078 |
|  | ROI Laterality*Volume*Paradigm*Process | 1 | 160.4 | 0.01 | 0.934 |
|  | ROI Laterality*Volume*Paradigm*Material | 2 | 160.4 | 0.38 | 0.682 |
|  | ROI Laterality*Volume*Process*Material | 2 | 160.4 | 0.36 | 0.696 |
|  | ROI Laterality*Paradigm*Process*Material | 2 | 160.4 | 0.26 | 0.770 |
|  | Volume*Paradigm*Process*Material | 2 | 160.4 | 2.10 | 0.126 |
|  | ROI Laterality*Volume*Paradigm*Process*Material | 2 | 160.4 | 0.81 | 0.446 |
| ERC | ROI Laterality | 1 | 166.61 | 0.08 | 0.777 |
|  | Volume | 1 | 41.30 | 0.80 | 0.376 |
|  | Paradigm | 1 | 159.97 | 1.21 | 0.274 |
|  | Process | 1 | 159.97 | 2.43 | 0.121 |
|  | Material | 2 | 159.97 | 3.60 | 0.029 |
|  | ROI Laterality*Volume | 1 | 167.91 | 0.17 | 0.677 |
|  | ROI Laterality*Paradigm | 1 | 159.97 | 0.09 | 0.766 |
|  | Volume*Paradigm | 1 | 159.97 | 2.16 | 0.143 |
|  | ROI Laterality*Process | 1 | 159.97 | 0.24 | 0.622 |
|  | Volume*Process | 1 | 159.97 | 0.24 | 0.628 |
|  | Paradigm*Process | 1 | 159.97 | 7.49 | 0.007 |
|  | ROI Laterality*Material | 2 | 159.97 | 0.06 | 0.942 |
|  | Volume*Material | 2 | 159.97 | 1.75 | 0.177 |
|  | Paradigm*Material | 2 | 159.97 | 0.09 | 0.915 |
|  | Process*Material | 2 | 159.97 | 1.07 | 0.346 |
|  | ROI Laterality*Volume*Paradigm | 1 | 159.97 | 0.21 | 0.648 |
|  | ROI Laterality*Volume*Process | 1 | 159.97 | 0.47 | 0.494 |
|  | ROI Laterality*Paradigm*Process | 1 | 159.97 | 0.01 | 0.917 |
|  | Volume*Paradigm*Process | 1 | 159.97 | 0.17 | 0.681 |
|  | ROI Laterality*Volume*Material | 2 | 159.97 | 0.09 | 0.911 |
|  | ROI Laterality*Paradigm*Material | 2 | 159.97 | 0.05 | 0.953 |
|  | Volume*Paradigm*Material | 2 | 159.97 | 0.60 | 0.549 |
|  | ROI Laterality*Process*Material | 2 | 159.97 | 0.59 | 0.554 |
|  | Volume*Process*Material | 2 | 159.97 | 0.59 | 0.554 |
|  | Paradigm*Process*Material | 2 | 159.97 | 0.82 | 0.443 |
|  | ROI Laterality*Volume*Paradigm*Process | 1 | 159.97 | 0.02 | 0.899 |
|  | ROI Laterality*Volume*Paradigm*Material | 2 | 159.97 | 0.09 | 0.918 |
|  | ROI Laterality*Volume*Process*Material | 2 | 159.97 | 1.13 | 0.325 |
|  | ROI Laterality*Paradigm*Process*Material | 2 | 159.97 | 0.14 | 0.868 |
|  | Volume*Paradigm*Process*Material | 2 | 159.97 | 0.06 | 0.945 |
|  | ROI Laterality*Volume*Paradigm*Process*Material | 2 | 159.97 | 0.25 | 0.779 |
| PHC | ROI Laterality | 1 | 167.99 | 0.07 | 0.797 |
|  | Volume | 1 | 37.35 | 1.99 | 0.167 |
|  | Paradigm | 1 | 159.08 | 1.40 | 0.238 |
|  | Process | 1 | 159.08 | 0.81 | 0.368 |
|  | Material | 2 | 159.08 | 10.92 | < 0.001 |
|  | ROI Laterality*Volume | 1 | 167.46 | 0.03 | 0.857 |
|  | ROI Laterality*Paradigm | 1 | 159.08 | 0.25 | 0.615 |
|  | Volume*Paradigm | 1 | 159.08 | 2.44 | 0.121 |
|  | ROI Laterality*Process | 1 | 159.08 | 0.13 | 0.720 |
|  | Volume*Process | 1 | 159.08 | 16.43 | < 0.001 |
|  | Paradigm*Process | 1 | 159.08 | 4.37 | 0.038 |
|  | ROI Laterality*Material | 2 | 159.08 | 0.07 | 0.928 |
|  | Volume*Material | 2 | 159.08 | 1.42 | 0.244 |
|  | Paradigm*Material | 2 | 159.08 | 0.54 | 0.586 |
|  | Process*Material | 2 | 159.08 | 0.50 | 0.610 |
|  | ROI Laterality*Volume*Paradigm | 1 | 159.08 | 0.33 | 0.564 |
|  | ROI Laterality*Volume*Process | 1 | 159.08 | 0.03 | 0.853 |
|  | ROI Laterality*Paradigm*Process | 1 | 159.08 | 0.74 | 0.391 |
|  | Volume*Paradigm*Process | 1 | 159.08 | 2.39 | 0.124 |
|  | ROI Laterality*Volume*Material | 2 | 159.08 | 0.08 | 0.920 |
|  | ROI Laterality*Paradigm*Material | 2 | 159.08 | 0.12 | 0.883 |
|  | Volume*Paradigm*Material | 2 | 159.08 | 0.03 | 0.970 |
|  | ROI Laterality*Process*Material | 2 | 159.08 | 1.22 | 0.297 |
|  | Volume*Process*Material | 2 | 159.08 | 2.06 | 0.131 |
|  | Paradigm*Process*Material | 2 | 159.08 | 1.15 | 0.320 |
|  | ROI Laterality*Volume*Paradigm*Process | 1 | 159.08 | 1.15 | 0.285 |
|  | ROI Laterality*Volume*Paradigm*Material | 2 | 159.08 | 0.23 | 0.798 |
|  | ROI Laterality*Volume*Process*Material | 2 | 159.08 | 2.08 | 0.128 |
|  | ROI Laterality*Paradigm*Process*Material | 2 | 159.08 | 0.07 | 0.930 |
|  | Volume*Paradigm*Process*Material | 2 | 159.08 | 2.69 | 0.071 |
|  | ROI Laterality*Volume*Paradigm*Process*Material | 2 | 159.08 | 0.17 | 0.847 |

Results from fitting a series of linear mixed-effects models, one for each volume of interest (HPC, ERC, PRC, PHC). Each model involved the following fixed effects and interactions thereof: ROI ROI Laterality (L, R); HPC/ERC/PRC/PHC volume; Paradigm (ROC, RDP); Process (Recollection, Familiarity); Material Type (Faces, Scenes, Words). **Key: PHC:** parahippocampal cortex; **PRC:** perirhinal cortex; **HPC:** hippocampus; **ERC:** entorhinal cortex; shaded cells: significant (p < 0.05) main effects and interactions of interest.

#### Supplementary figure 5

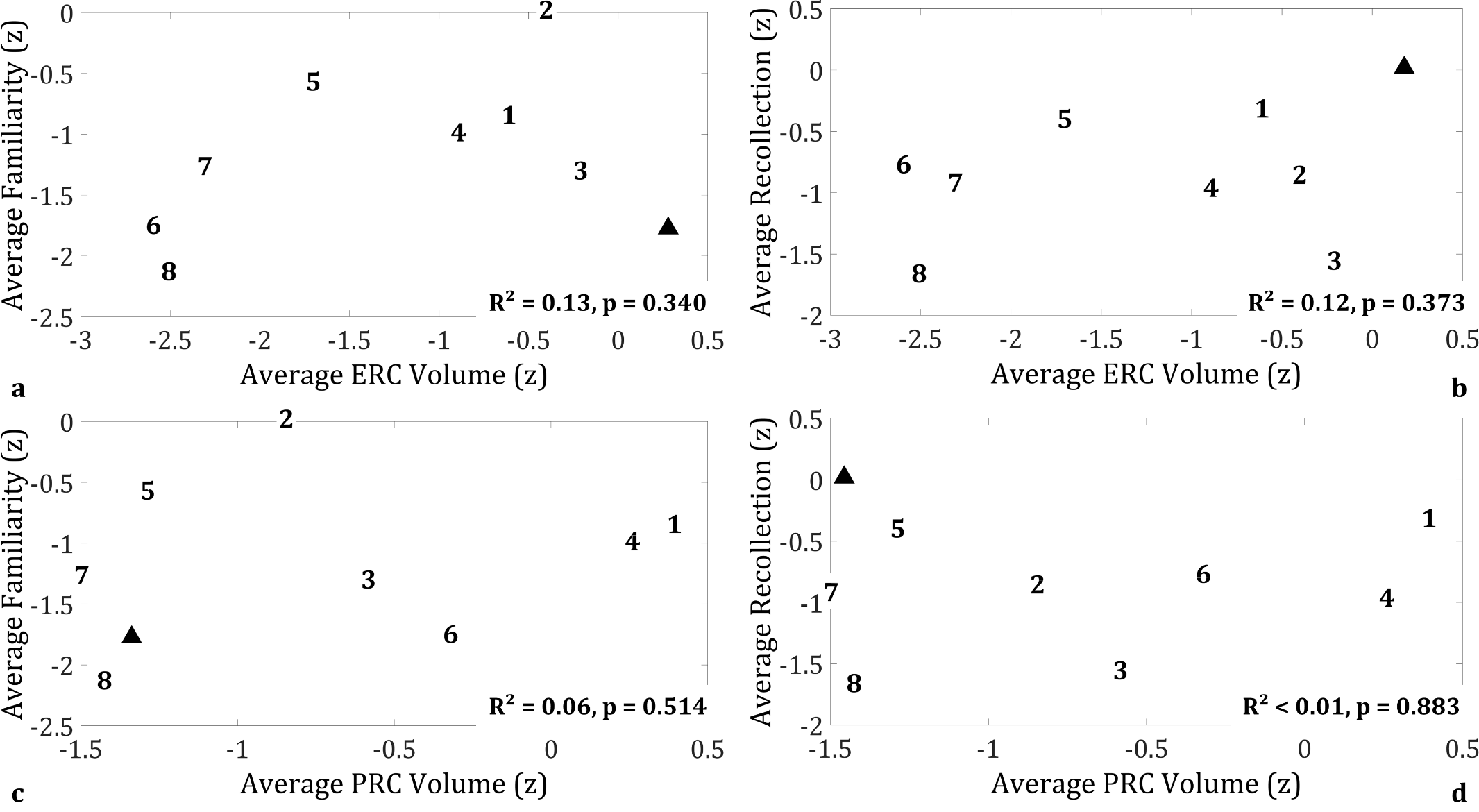

Brain-behaviour relationships between ERC/PRC volumes and familiarity/recollection process strength across patients; no relationships were identified between these volumes and recollection/familiarity across patients; **key: ERC:** entorhinal cortex; **PRC:** perirhinal cortex; **z:** volumes are expressed as z-scores, based on the mean and standard deviation of the volumes of the 48 healthy controls whose MTL structures were manually delineated (see [(Argyropoulos et al., 2019)] for details); familiarity and recollection estimates are expressed as z-scores, based on the mean and standard deviation of the 14 CTRs that completed the two tasks.
